# Light and sex modify *Snord116* genotype effects on metabolism, behavior, and imprinted gene networks following circadian entrainment

**DOI:** 10.1101/2025.05.01.651733

**Authors:** Aron Judd P. Mendiola, Jennifer M. Rutkowsky, Kari Neier, Sophia Hakam, Osman Sharifi, Callum Donnelly, Christina Torres, Cielo Hernandez, Dag H. Yasui, Jon J. Ramsey, Janine M. LaSalle

## Abstract

Mammals utilize imprinted and X-linked epigenetic mechanisms in development, metabolism, and behavior. Imprinted genes, including Prader-Willi syndrome *Snord116* noncoding RNAs, are implicated in the regulation of sleep and circadian rhythms through poorly understood mechanisms. Utilizing mouse models of *Snord116* deficiency and overexpression, we performed an integrated, sex-stratified analysis of free running behaviors, indirect calorimetry, and cortical transcriptomes following entrainment to a 22 hour light:dark T-cycle. We observed significant interactions of sex, entrainment, and *Snord116* genotype in period length at baseline and after-effects of post-entrainment. *Snord116* deletion’s effect on respiratory exchange ratio was light sensitive, with sex and entrainment effects dominant under total darkness. *Snord116* genotype impacted both rhythmic and non-rhythmic cortical gene networks that integrated sex, light, and entrainment effects with genotype-phenotype correlations. A co-expressed gene network enriched for imprinted, *Snord116*-target, and *Xist*-proximal long noncoding RNAs was identified as a light-sensitive regulatory hub of sexual dimorphic responses to a dynamic environment.

## Introduction

Daily and seasonal cycles of light, temperature, and feeding are environmental and metabolic inputs that play an important role in the synchronization of the core circadian clock with the rhythmic patterns of sleep as well as many physiological and behavioral processes in peripheral tissues (Blasiak et al., 2017; Legates et al., 2014; Mukherji et al., 2015; Wright et al., 2013). The genetically encoded circadian clock and the environmentally regulated diurnal cycle are integrated by a complex regulatory feedback network which acts at the chromatin, transcriptional, and translational levels to coordinate biological and environmental rhythms (Koike et al., 2012; Papazyan et al., 2016; Takahashi, 2017). In mammals, the core circadian clock resides in the suprachiasmatic nucleus of the hypothalamus; however, almost half of all transcripts, both protein-coding and non-coding, exhibit diurnal rhythms in one or more peripheral tissues (Yan et al., 2008; Zhang et al., 2014). The cerebral cortex utilizes more than 20% of the body’s energy (Herculano-Houzel, 2011), influencing both metabolism and behavior, but the cortex’s involvement in circadian entrainment of metabolism and behavior to a dynamic environment is poorly understood.

Sleep disturbances are common in human neurodevelopmental disorders associated with the 15q11-q13 imprinted locus (Esbensen and Schwichtenberg, 2016) and their mouse models (Lassi et al., 2016; Tucci, 2016). This includes Prader-Willi syndrome (PWS) and Angelman syndrome (AS), which are characterized by excessive daytime sleepiness and shorter sleep duration, respectively (Colas et al., 2005; Gibbs et al., 2013; Lassi et al., 2016; Thibert et al., 2013). PWS is characterized by a failure to thrive and hypotonia in infancy, followed by hyperphagia as well as metabolic, behavioral, cognitive, and sleep abnormalities (Butler et al., 2002; Cassidy et al., 2012). Small paternal deletions have defined the minimal PWS critical region to encompass *SNORD116* (Bieth et al., 2015; de Smith et al., 2009; Duker et al., 2010; Sahoo et al., 2008). The paternally expressed *SNORD116* locus includes repeats of small nucleolar RNAs (snoRNAs) processed from a long polycistronic transcript. The spliced host gene of *SNORD116* (*116HG/Snhg14*) forms a nuclear long noncoding RNA (lncRNA) “cloud” that localizes to its site of transcription and is diurnally regulated in size (Powell et al., 2013a; Powell et al., 2013b). Other lncRNA forming clouds at their own chromosomal loci include female specific *Xist*, and other imprinted locus genes *Meg3* and *Kcnq1ot1* (Burnett et al., 2017; Ding et al., 2008; Powell et al., 2013a; Powell et al., 2013b; Qian et al., 2016). Our prior studies of male mice carrying a ∼150kb deletion of *Snord116* (*Snord116^+/-^*) revealed a role of this locus in the diurnal regulation of genes with circadian, metabolic, and epigenetic functions, including “circadian entrainment” as the top enriched pathway (Coulson et al., 2018b).

To test the hypothesis that *Snord116* orchestrates diurnal rhythms of behavior, metabolism, and cortical gene expression in response to a dynamic lighting environment, we performed a three-week T-cycle entrainment to 11:11 versus 12:12 light:dark cycles. We utilized transgenic mice of different *Snord116* copy number, following entrainment with sex-stratified and integrated analysis of free running behavior, metabolism, and transcriptomics under different lighting conditions. We observed that multiple phenotypes associated with *Snord116* genotypes were modified by both sex and external light cues. Females specifically retained significant transcriptional and metabolic effects of prior entrainment in constant darkness that correlated with *Xist*-associated networks of genes, including imprinted lncRNAs. Together, these results are impactful in understanding how transcriptional sex differences can modify a gene-environment interaction at an imprinted gene locus relevant to human cognition and metabolism.

## Results

### Experimental design integrates Snord116 genotype, lighting, sex, and circadian entrainment with phenotypes of free running behavior, metabolism, and cortical transcriptomes

We designed a multidimensional study to test the hypothesis that *Snord116* regulates rhythmic gene expression patterns in response to a changing environment using two genetic mouse models with different copy numbers of *Snord116* (Coulson et al., 2018a; Ding et al., 2005). The four genotypes of mice include *Snord116*^+/+^*Ctg*^-/-^ (WT/WT; Wild Type), *Snord116*^+/+^*Ctg*^+/-^ (WT/TG; Overexpression), *Snord116*^+/-^*Ctg*^-/-^ (HET/WT; Deletion), and *Snord116*^+/-^*Ctg*^+/-^ (HET/TG; Compensation) with fewer HET/TG than WT/TG than expected, but otherwise expected ratios (Data S1). Both males and females were included to investigate sex-specific responses to genotype and experimental lighting conditions (Figure 1). Each genotype and sex of mice were born and housed under controlled lighting conditions of 12 h light and 12 h dark (12:12 L:D). At 8 weeks of age, the mice were divided and entrained for 3 weeks to two different lighting conditions: either 12:12 L:D control or 11h light and 11 h dark (11:11 L:D) entrainment treatments. The mice were split into two phenotyping groups to determine the effects of entrainment, sex, and genotype on diurnal free running activity and metabolism in different lighting conditions. In the first group, voluntary running wheel measurements began at the start of the third week of entrainment, followed by a reversion back to 12:12 L:D to study the after-effects of returning 11:11 entrained mice to 12:12 conditions. In the second group, metabolic measurements using indirect calorimetry (CLAMS) began after 22 days of entrainment, including respiratory exchange rate (RER), heat (energy expenditure, kcal), X-tot (activity derived from horizontal beam breaks), energy intake (kcal/h feeding), and sleep (inactivity derived from the lack of beam breaks). After 24 h of acclimation to chambers, metabolic measurements were first collected under entrained lighting conditions (L:D) for 2 days, followed by 4 days of additional measurements in constant darkness (D:D). All metabolic data were divided into light hour and dark hour measurements, including the D:D conditions that were divided by prior light and dark hours. At the end of each study, mouse brains were harvested at zeitgeber time 6 (ZT6), that we had previously determined as the critical time point of epigenetic and transcriptional effects of *Snord116* in the prefrontal cortex (Coulson et al., 2018b; Powell et al., 2013a). To determine transcriptional differences at the interface of genotype, entrainment, sex, RNAseq was performed. In addition, a cortical circadian transcriptome atlas was generated from a separate cohort of wild-type age-and sex-matched mice collected every 3 hours in 12:12 LD home cage conditions to determine transcriptional rhythmicity. A consensus network analysis was used to integrate transcriptional networks with behavioral and metabolic phenotypes using weighted gene correlation network analysis (WGCNA) (Langfelder and Horvath, 2008). These experiments were designed to test how loss of endogenous *Snord116* and the gain of transgenic *Snord116* impact multiple phenotypes of behavior, metabolism, and cortical transcription in response to changing environmental conditions of light and dark.

**Figure 1.**
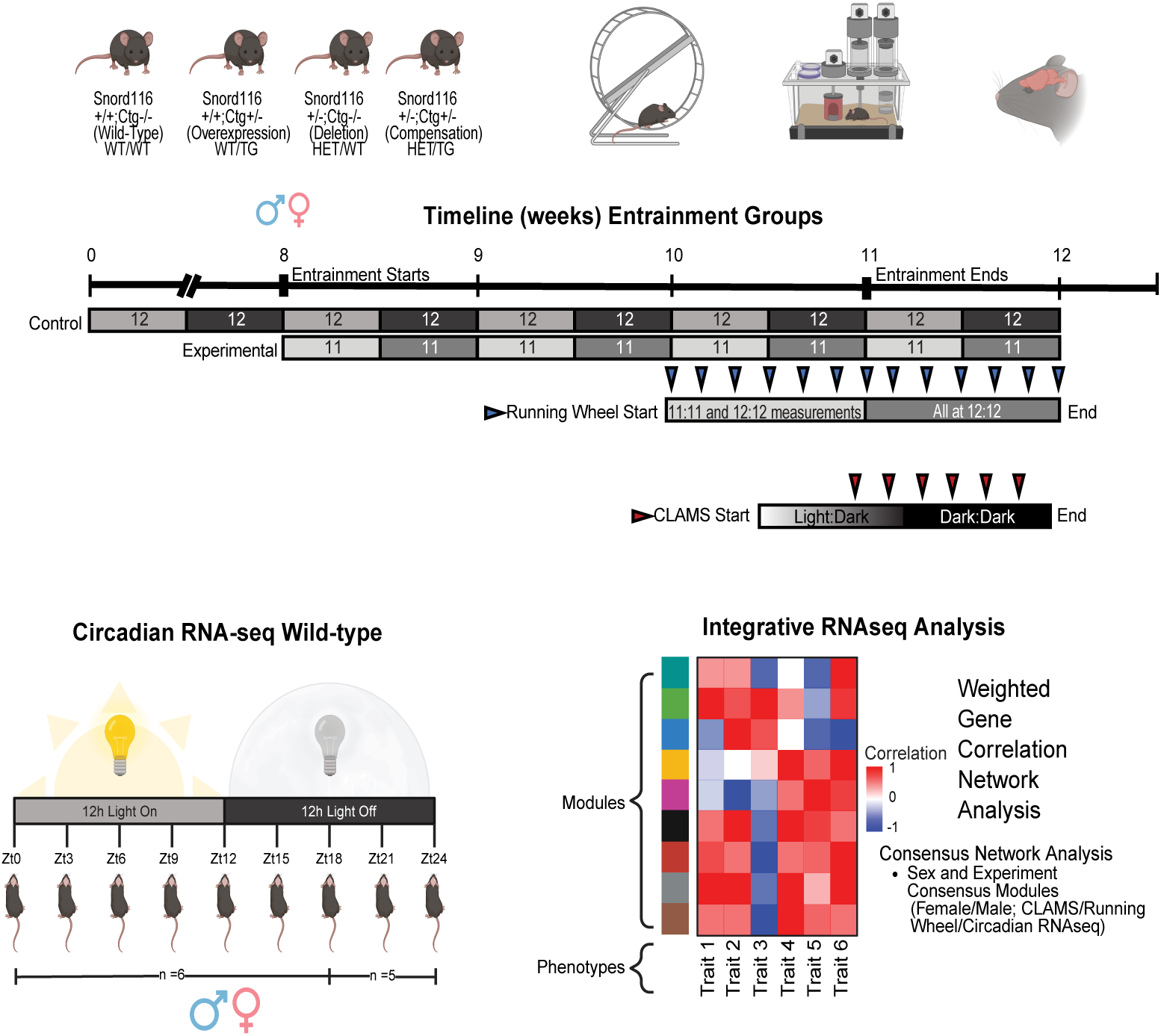
Experimental design to identify the effects of *Snord116* copy number and circadian entrainment on behavior, metabolism, and gene networks in the prefrontal cortex. Four *Snord116* mouse model genotypes of both sexes were entrained to either control 12h light and 12h dark (12:12) or experimental 11h light and 11h dark (11:11). Running wheel (L:D) collections were split into week 1 and week 2 measurements. At week 2, 11:11 entrainment group was returned to 12:12 lighting conditions. Indirect calorimetry/CLAMS (D:D) measurements were collected under both normal light-dark (L:D) and constant darkness (D:D) lighting conditions. At the end of each study, mouse brain cortex samples were collected at the peak of the original light phase (ZT6) for RNAseq experiments. A separate group of circadian RNA-seq wild-type mice were included in a consensus network analysis of prefrontal cortex, with mouse cortex collected every 3 hours over 24 h 12:12 L:D home cage conditions. Integration of RNAseq analysis with phenotype measurements and experimental conditions was performed with weighted gene correlation network analysis (WGCNA).

### Significant interactions of sex, entrainment, and Snord116 genotype were observed in period, amplitude, and activity before or after circadian entrainment

To characterize the effects of circadian entrainment on free running behavior in *Snord116* deletion and overexpressing mouse models, period, amplitude, total running distance, and total running time were measured. 2-way ANOVA statistics of entrainment and genotype effects identified significant effects for entrainment and genotype for multiple measurements but there were many sex differences in these effects (Figure 2A and Data S2), so we also performed 3-way ANOVA tests to include sex (Figure 2B-C). Period measurements demonstrated that while all mice adapted to entrained lighting conditions, seen as shorter period for 11:11 than 12:12 entrainment, there were significant interaction effects with sex and genotype in period lengths at baseline and “after-effects” in the first 4 days after returning to 12:12 (Figure 2B). Overall, females showed significantly higher amplitude, total time running, and total distance running during week 1 (base) compared to males regardless of genotype or entrainment (Figure 2C and Figure S1), as expected from prior studies (Bronstein et al., 1975; Fischer et al., 2016; Lightfoot, 2008; Lightfoot et al., 2004; Perrigo and Bronson, 1985; Rosenfeld, 2017). Interestingly, the most significant genotype effects in free running behavior were female-specific, include significantly lower baseline period, amplitude (both weeks), and total time and distance running in *Snord116* deletion compared to wild-type females (Figure 2A, Figure S1, Data S2). However, males showed significant interactions between entrainment and genotype for base period and first 4-day after-effects, where 11:11 HET/WT and HET/TG mice had a longer period at base and returned to a 24 h T-cycle sooner than mutant females (Figure 2A-B). There were also significant genotype and sex effects on total distance and running time, with males and *Snord116* deletion mice showing lower activity during and after entrainment (Figure S1).

**Figure 2.**
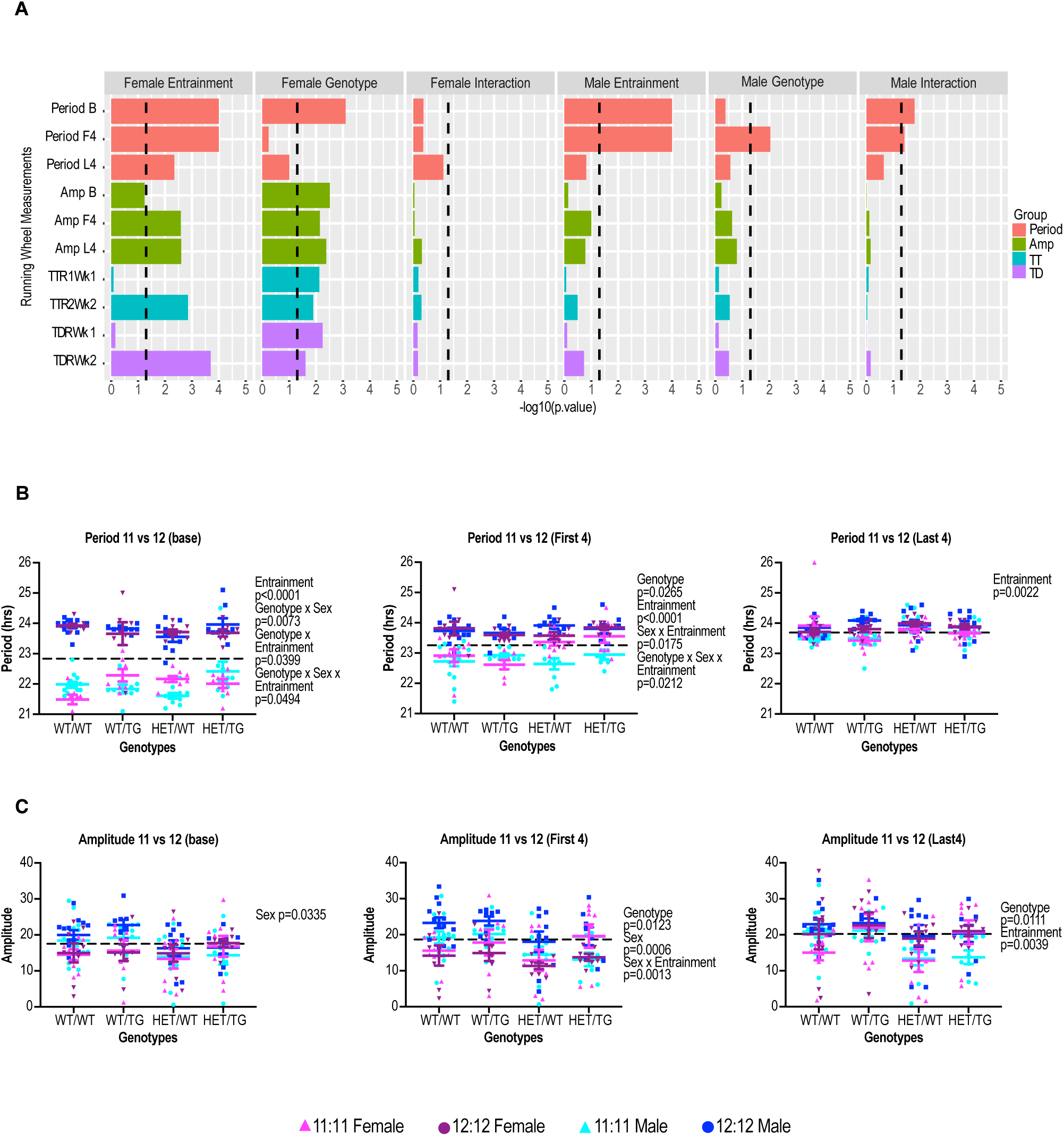
*Snord116* genotype interacted with both sex and entrainment effects in free running behavior. A) Running wheel 2-way ANOVA statistics of entrainment, genotype, and interaction effects. Multiple pairwise comparisons of different genotypes were also performed with Fisher’s LSD test (Data S2-S3). Amp, amplitude; TTR, total time running; TDR, total distance running; B, Baseline; F4, first four days; L4, last four days; Wk, week. B) Scatter plots of running period length measurements for each mouse by genotype and sex, taken at baseline, week 1 (entrainment), and during the first and last four days of week 2 (after-effects) with 3-way ANOVA statistics of entrainment, genotype, and sex. C) Scatter plots of running wheel amplitude measurements taken for each genotype and sex at baseline, week 1, and during the first and last four days of week 2, with 3-way ANOVA statistics for entrainment, genotype, and sex.

Interestingly, sex effects were no longer significant by 3-way ANOVA during the last 4 days of free running measurements under 12:12 conditions that were included to examine the lasting behavioral adaptations to novel lighting conditions (Figure 2). However, by 2-way ANOVA, there were female-specific lasting effects of the combined effects of HET genotypes and 11:11 entrainment and on period, amplitude, running time, and running distance (Data S2). To determine if *Snord116* deletion effects were rescued in the compensation model (HET/TG), we required the following statements to be true: 1) deletion must be significantly different from wild-type, 2) compensation must be greater than or equal to wild-type, 3) compensation must be significantly different from the deletion in the same direction as the wild-type to *Snord116* deletion comparison. By these criteria, none of the *Snord116* deletion behavioral effects were compensated by the *Snord116* transgene (Data S3). Together these results support the requirement of endogenous *Snord116* for behavioral adaptations of circadian entrainment but also show that transgenic *Snord116* copies expressed from a separate chromosomal locus are insufficient to rescue free running phenotypes. In addition, these behavioral effects of *Snord116* deficiency following circadian entrainment were predominantly female-specific.

### Sex and lighting conditions impacted Snord116 genotype effects on metabolic measurements following circadian entrainment

To determine the metabolic effect of circadian entrainment and *Snord116* deficiency and/or overexpression, mice were monitored in indirect calorimetry chambers continuously during light and dark hours and under both L:D and D:D conditions. 2-way ANOVA statistics of entrainment and genotype effects identified significant effects for entrainment, genotype, and their interaction for multiple measurements under different lighting conditions (Figure 3A, Data S4). Sex differences were also further investigated by 3-way ANOVA (Figure 3B-C). As expected, based on prior analyses of male *Snord116* deletion housed in 12:12 L:D conditions (Powell et al., 2013a), HET/WT mice had significantly reduced respiratory exchange ratio (RER) during light hours compared to WT/WT in both sexes, indicating elevated use of lipids compared to carbohydrates as energy. This effect was not rescued in the HET/TG compensation model (Figure 3A-B). Specifically for females in L:D, there was also a significant interaction between 11:11 entrainment and genotype for RER during dark hours. But these genotype effects on RER were light sensitive, as genotype effects on RER were not present under constant darkness. In contrast, mice in constant darkness showed significant lasting effects of 11:11 entrainment in both sexes, which was observed in RER, energy expenditure, energy intake, horizontal activity, and sleep (Figure 3A, Data S4). But 11:11 entrained male mice were distinct in showing the lowest RER and activity levels that were most apparent under constant darkness (Figure 3B-C). Similar to what was observed in free running activity, light hour horizontal activity (Xtot) in L:D was lower in males and HET/WT mice in L:D, but not compensated by HET/TG.

**Figure 3.**
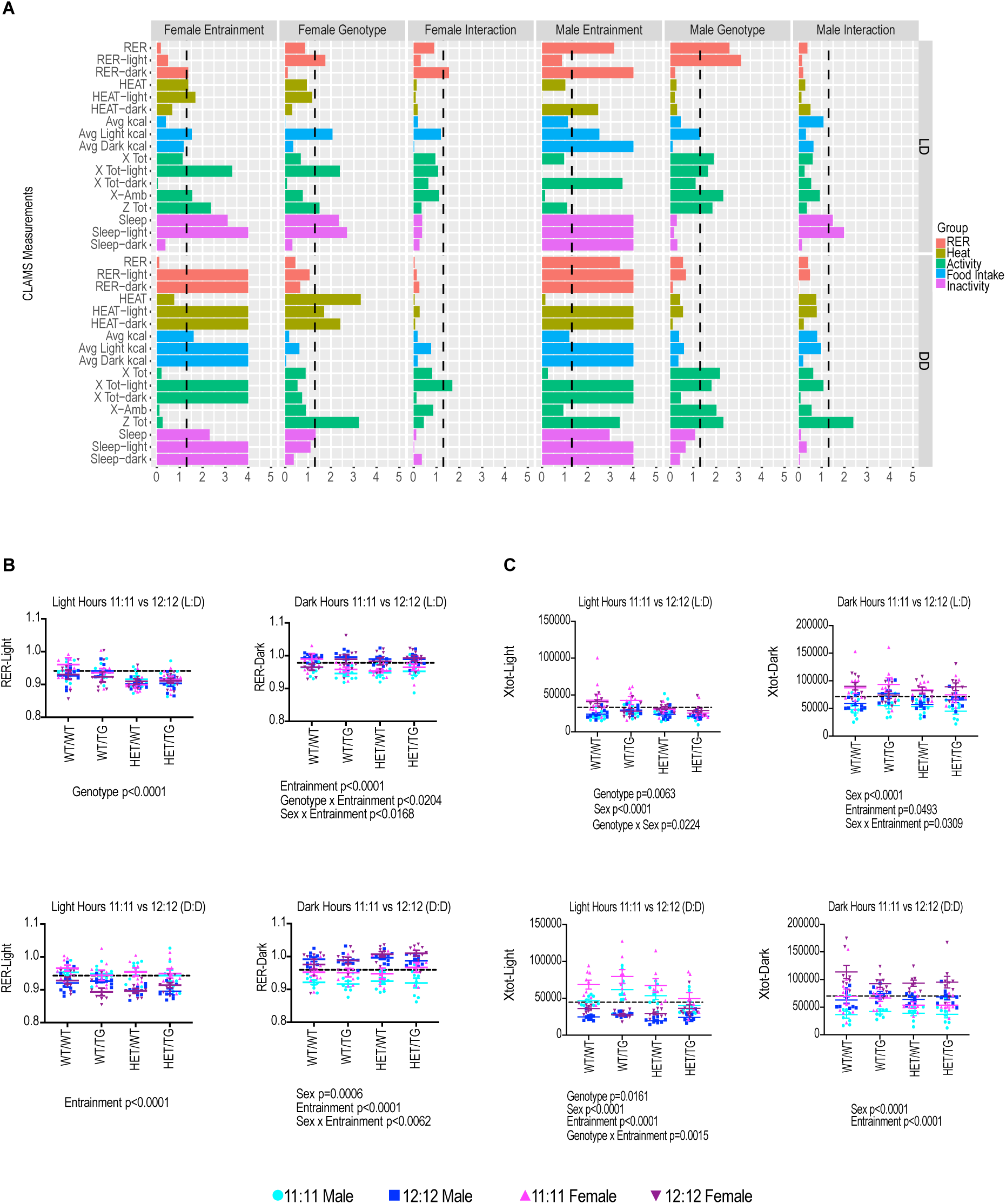
*Snord116* genotype effects on lipid oxidation during sleep (light RER) were dependent on external light cues independent of sex or entrainment, while activity levels showed multiple interactions. (A) Indirect calorimetry (CLAMS) 2-way ANOVA statistics of entrainment, genotype, and interaction effects separated by normal light-dark (L:D) and constant darkness (D:D) lighting conditions. Non-corrected genotype multiple pairwise comparisons with Fisher’s LSD test was also performed. Data S4 has a full list of p-values. (B) Scatter plots of RER measurements for each genotype and sex taken during light hours and dark hours of L:D and D:D lighting conditions with 3-way ANOVA statistics of entrainment, genotype, and sex. (C) Scatter plots of activity (X-tot) measurements for each genotype and sex taken during light hours and dark hours of L:D and D:D lighting conditions with 3-way ANOVA statistics of entrainment, genotype, and sex.

These results are consistent with the after-effects of entrainment being strongest in D:D conditions since the absence of external light cues cause mice to use their internal homeostatic cues to maintain rhythms of behavior and metabolism. But these results also show that female and male mice have distinct metabolic responses to 11:11 entrainment and *Snord116* genotype. Interestingly, female mice showed *Snord116* genotype effects of decreased energy expenditure specifically in D:D compared to L:D, while male mice maintained genotype differences in their activity under both lighting conditions (Figure S2). Both sexes show expected genotype (but not entrainment) significant effects of reduced body weight, % fat, fat mass, and lean mass in *Snord116* deletion mice that were not reversed in compensation mice (Figure S3, Data S5).

Overall, female mice were more impacted by *Snord116* genotype than males in their energy expenditure under D:D, as well as energy intake (kcal/h) and sleep phenotypes under L:D (Figure 3A, Figures S3-S4). These results indicate that endogenous *Snord116* is involved in coordinating metabolic adaptations in response to changing lighting conditions, with more profound effects in females.

### Snord116 genotypes and phenotypes correlated with both rhythmic and nonrhythmic cortical gene networks but correlations differed by sex and lighting condition

To test the hypothesis that *Snord116* is involved in regulation of rhythmic gene networks in the prefrontal cortex that can be regulating phenotypic differences, we used blockwise consensus WGCNA to combine transcriptomic data sets from cortical RNA collected from circadian atlas (WT mice in home cage, 12:12, L:D, ZT0-24), experimental L:D ZT6 (running wheel), and experimental D:D ZT6 (CLAMS) experiments. The consensus modules approach allowed us to compare the same gene networks across sexes and samples collected under different experimental conditions. WGCNA generates module gene networks based on similarity of gene expression through midweight correlation statistics. The consensus networks produced universal modules that are highly preserved across sex, experiment, and all other conditions (Figure S5, Data S6). RAIN circadian statistics was used to identify phases and rhythmicity of module network eigenvalues. 14 significantly rhythmic female modules and 17 significantly rhythmic male modules were identified, indicating sex-specific differences in rhythmic gene expression in cortex (Figure 4A). The expected rhythmic expression patterns of core clock genes were observed in both sexes (Figure 4B, 4C). *Clock*, *Arntl*, *Cry1*, and *Cry2* were found in the largest grey “catch-all” module, while *Per1* and *Per2* were found in the light cyan and midnight blue modules, respectively. *Per1* and *Per2* both belong to significantly rhythmic modules with a shared phase of ZT18, indicating that they are in networks with other genes that are highly expressed at that timepoint. To track expression of *Snord116*, we used *Snhg14* and *Snrpn* transcripts. *Snhg14* and *Snrpn* were found in the statistically non-rhythmic grey module, but their peak phase was distinct between females (ZT12) and males (ZT6) (Fig 4B-C). These results suggested that a subset of gene networks regulated by *Snord116* and circadian entrainment may be different between sexes in the prefrontal cortex.

**Figure 4.**
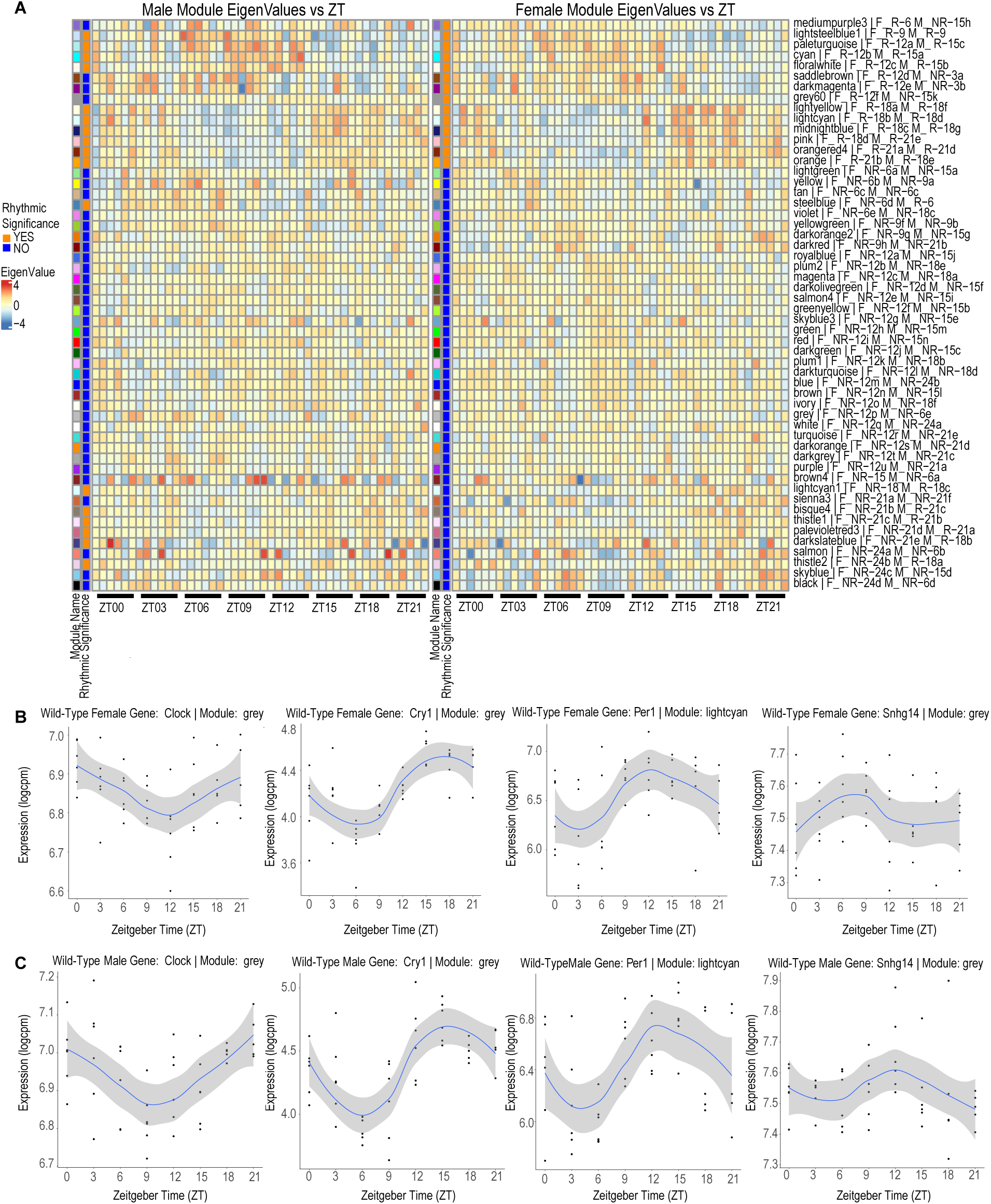
A sex-stratified rhythmic co-expression gene network atlas of the prefrontal mouse cortex was defined by blockwise consensus WGCNA. (A) Heatmaps of rhythmic coexpression gene networks identified by RAIN circadian statistics of male (left) and female (right) mouse prefrontal cortex. The module eigenvalues are plotted against the ZT to show the phase of each module network and are ordered based on rhythmic significance of female modules. Modules are given arbitrary color names, but additional names are given to reflect rhythmicity for each sex: F, female; M, male; R, rhythmic; NR, nonrhythmic; significant (orange), non-significant (blue), and the number represents the peak phase in ZT. (B) Loess plots of core clock genes (*Clock, Cry1, Cry2, Per1, Per2*) expression in female mice. (C) Loess plots of core clock genes expression in male mice and includes Loess plots of *Snhg14* and *Snrpn*.

To connect the phenotypic measurements collected from behavioral and metabolic experiments to transcriptional changes caused by genotype and entrainment, WGCNA module-trait correlations were performed. In the four heatmaps summarizing all experimental correlations, female mice housed in constant darkness (D:D) showed the most significant transcriptional correlations to entrainment, genotype, and phenotype measurements compared to D:D males and L:D mice of both sexes (Figure 5). In D:D females, 20 gene modules were highly significantly correlated (p < 0.001) with 11:11 entrainment (14 positively and 6 negatively, red versus blue, respectively) compared to only 2 in D:D males (Figure 5A, 5C). In females under constant darkness, gene modules with high correlations to entrainment also showed strong associations with RER, heat, activity, feeding, and sleep. Interestingly, modules that were positively correlated with 11:11 entrainment were also positively associated (red) with prior light hour measurements but negatively associated (blue) with prior dark hour measurements, and vice-versa. For females under constant darkness conditions, 10 modules showed lower expression in HET/WT versus WT/WT, and 9 of these also met criteria for compensation by the *Snord116* transgene described above. These 9 compensation-sensitive modules included three rhythmic (paleturquoise, cyan, floralwhite) modules with a sex-specific peak at ZT12 in females, compared to ZT15 in males, as well as 4 nonrhythmic (darkorange2, darkolivegeen, red, brown, white), and one module that was rhythmic in females but not males (grey60) (Figure 5A). Many additional gene modules were significantly correlated in opposite directions between the compensation versus deletion and the overexpression versus compensation comparisons, suggesting that the transgene’s effect on transcriptional changes was stronger in the *Snord116* deletion than WT background.

**Figure 5.**
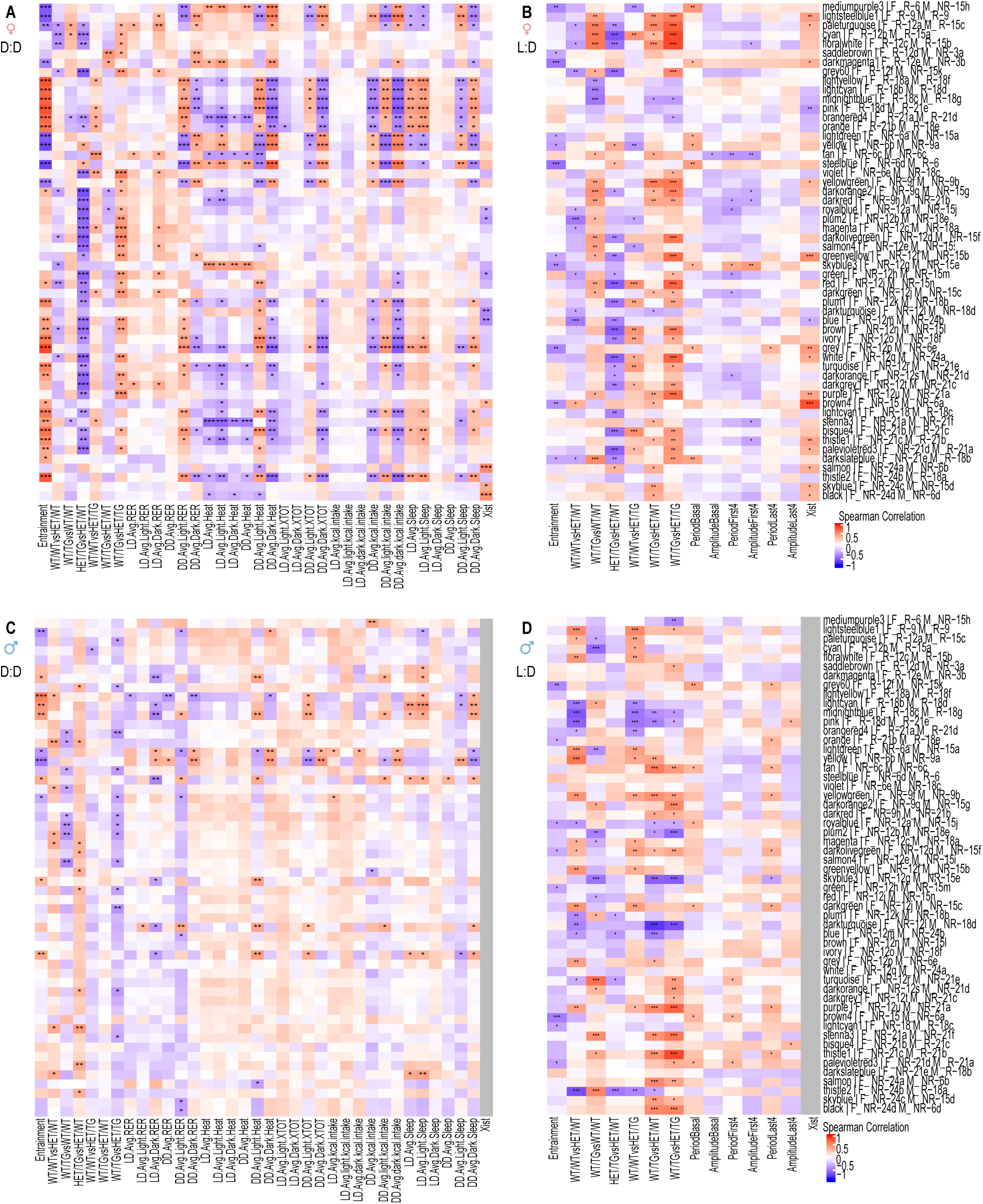
Sex-specific cortical gene networks connect *Snord116* compensatory effects on transcription with metabolic phenotypes. All consensus WGCNA modules were labeled and ordered on the y-axes according to those defined in Figure 4. (A) D:D female module-trait correlation heatmap with genotype and indirect calorimetry phenotypes. (B) L:D female module-trait correlation heatmap with genotype and running wheel phenotypes. (C) D:D male module-trait correlation heatmap. (D) L:D male module-trait correlation heatmap. Colors represent positive (red > 0) and negative (blue < 0) correlations and significance (***) p-value < 0.001, (**) p-value < 0.01, and (*) p-value < 0.05 by Spearman correlation. Genotype comparisons were pair-wise, with highest *Snord116* copy number listed first and coded as baseline, as shown on the x-axis labels.

Unlike under constant darkness, both sexes showed significant correlations with entrainment and/or *Snord116* genotype in diurnal L:D conditions, although sex differences were observed in the direction of these effects and the significance of specific modules (Figure 5B and 5D). For females under L:D conditions, 11 modules were significantly associated in WT/WT versus HET/WT genotype, with all these modules showing lower expression in *Snord116* deletion cortex except one (brown4) module, and 6 of these modules meeting criteria for being compensation-sensitive (pale turquoise, floralwhite, grey60, green, blue, darkslateblue) (Figure 5B). In contrast, although 22 modules were positively or negatively associated with WT/WT versus HET/WT genotype in L:D males, only three (plum1, blue, turquoise) modules were compensation-sensitive (Figure 5D). These results indicate that the exogenous *Snord116* transgene was able to compensate for some sex-differential transcriptional changes in the endogenous *Snord116* deletion cortex even though the behavioral and metabolic phenotypes were not significantly rescued.

To further characterize functions of the genes within each module, we performed an analysis of gene functions through Gene Ontology (GO) terms, gene pathways through KEGG, and human phenotypes through database of Genotypes and Phenotyes (dbGaP)(Data S7-S11). DbGaP terms were enriched for several relevant human metabolic phenotypes in genotype/phenotype correlated modules, including body weight (turquoise: *CNTNAP2, NBEA, USP34, NRIP1, CSMD1, ASTN1*; red: *PTPRD, SLC24A2, PLCB4, KCNH7, ATRNL1, CNTN3, CTNNA2*), adiposity (floralwhite: *FTO*) bone density (brown: *THRB, SEMA3C, DNAJC6, ERBB4*), and sex hormones (darkorange2: *GUCY1A2*; ivory: *IRS1*) (Figure S6). The top 100 KEGG terms that were significantly enriched within genotype/phenotype correlated modules included circadian entrainment (darkorange2: *GUCY1A2, PRKCB, PLCB1*; blue: *ADCY9, ADCY2, CAMK2G, GRIN1;* darkgrey: *PRKCG, GNG2, GNAQ*; salmon: *GNG7, GNB4, RYR3, ADCY5*), thermogenesis (magenta: *NDUFA9, CREB3, ATP5B, KDM1A, ATP5A1, UQCRC1, NDUFV1*), glycolysis/gluconeogenesis (magenta: *MINPP1, PFKL, TPI1, PGAM1, DLAT*), thyroid hormone (darkorange2: *GSK3B, PRKCB, PLCB1*; blue: *ADCY9, ATP1A3, ADCY2, ATP1A1*), and other functional pathways that are relevant to PWS. Furthermore, enrichment analysis these modules of other RNAseq datasets (RNADD, Data S12) showed significant enrichments to our previously published analyses of this PWS mouse model (GSE43575), as expected.

Some gene modules that differed in diurnal rhythmicity between males and females (Figures 4-5) also showed functional enrichments for sexually dimorphic traits or diseases (Figure S6). For example, mediumpurple3 was rhythmic in females but not males, showed female-specific decreased expression with 11:11 entrainment, and was significantly enriched for KEGG terms “menopause” and “menarche.” In contrast, the lightcyan1 module was rhythmic in males but not females, showed male-specific decreased expression following 11:11 entrainment in L:D, and was enriched for dbGAP traits of abdominal fat, HDL cholesterol, and head and neck neoplasms (3:1 M:F ratio). These sexually dimorphic rhythmic gene modules are therefore potentially useful in better understanding sex differences relevant to clinical traits and diseases.

### Female-specific Xist transcript levels correlated with modules showing female-specific genotype and phenotype effects, including a network of imprinted and Xist-colocalized lncRNAs

Because of the strong sex differences observed in phenotypes, rhythmic gene modules, and module-trait correlations, we also explored the hypothesis that the female-specific noncoding gene *Xist* may correlate with modules showing female-specific module-trait correlations. We therefore added *Xist* transcript levels as a trait to the correlation analyses shown in Figure 5. This is because *Xist* was not included within the universal module gene networks that were selected to work across both sexes. For female cortex collected under constant darkness, 9 non-rhythmic genes modules were significantly associated with *Xist* transcript levels (Figure 5A). The female-specific *Xist* transcript was associated with 16 gene modules under L:D conditions, including 3 (paleturquoise, blue, brown4) that were also significantly changed by *Snord116* deletion (Figure 5B).

Interestingly, the imprinted noncoding *Meg3* lncRNA was the top hub gene for module brown4 (Data S6). The *Meg3* gene DNA locus was previously reported to be a direct target of *Snord116* host gene binding and *Meg3* transcript was highly upregulated at ZT6 but not ZT12 in *Snord116* deletion cortex compared to WT (Coulson et al., 2018b). Consistent with our prior *Meg3* results, brown4 transcript levels were elevated in L:D *Snord116* deletion females, as well as highly correlated with *Xist* (Figures 5B). In females, entrainment effects on brown4 transcript levels were in opposite directions depending on lighting conditions, significantly higher in 11:11 compared to 12:12 treatment under D:D but lower under L:D conditions, suggesting that females have a distinct mechanism regulating the brown4 network in the absence of light cues, explored in Sankey plots (Figure 6). In D:D conditions, brown4 transcript levels lost correlation with *Snord116* genotype compared to L:D in females but maintained positive correlation with *Xist.* In the absence of light cues (D:D), two rhythmic modules, lightsteelblue1 and paleturquoise, showed multiple correlations with female metabolic phenotypes, including RER, heat, energy intake, and sleep (Figures 5A, 6). While no genotype correlations were observed for brown4 in males, brown4 transcript levels in L:D male cortex positively correlated with base and first 4-day period, phenotypes that were *Snord116* deletion sensitive (Figures 2B, 5D).

**Figure 6.**
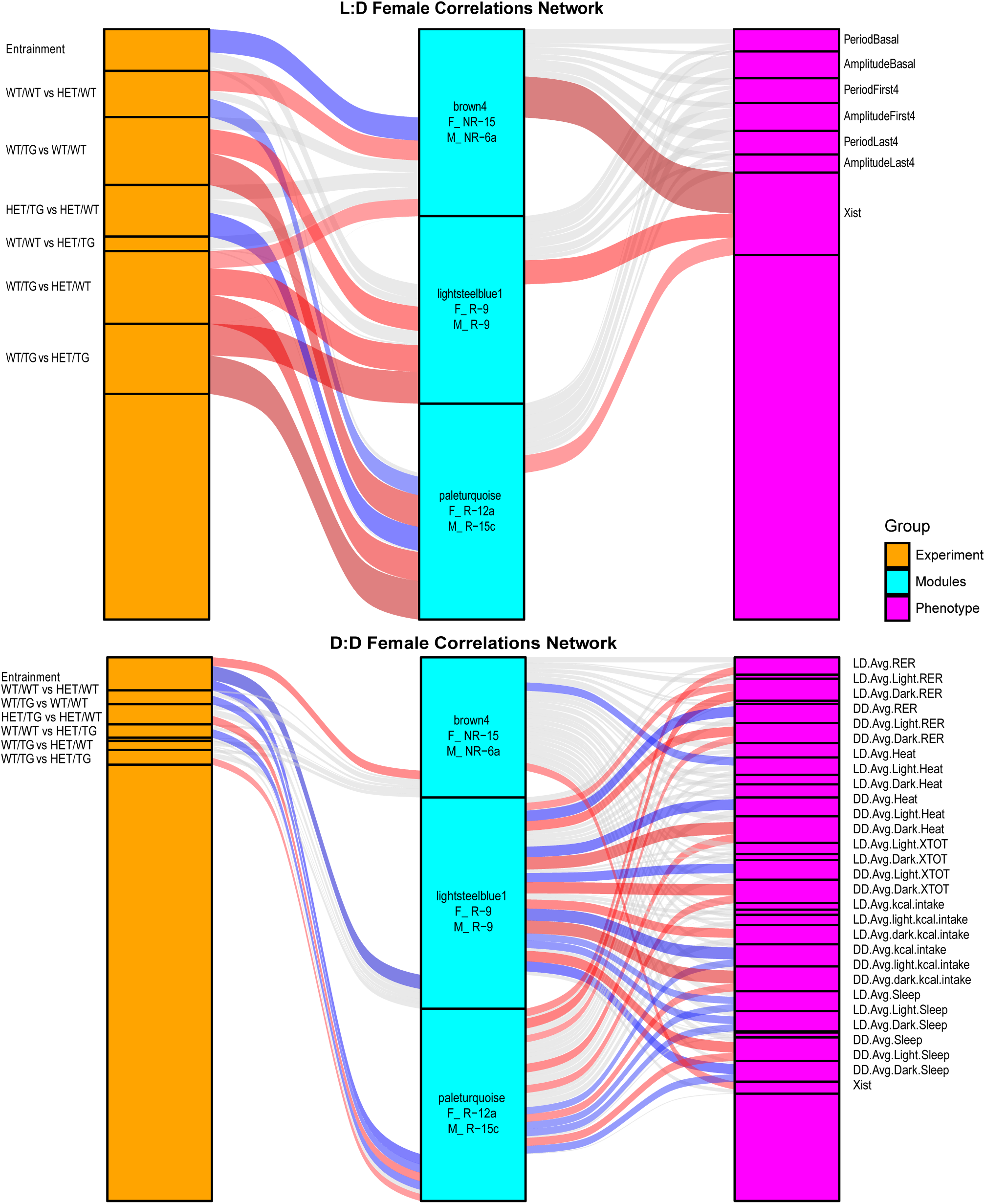
Sankey diagrams of genotypes, transcriptional modules, and phenotypes correlated with *Xist* in females showing dynamic changes in L:D versus D:D. Experimental conditions (genotypes or entrainment) are on the left, three highly dynamic correlated transcriptional modules are in the middle, and phenotypes including *Xist* are on the right columns. Overall, correlations between *Snord116* genotypes, *Xist*, and these transcription gene networks were stronger in L:D conditions (top), while correlations between these same networks with metabolic and sleep phenotypes were stronger in the absence of external light cues (D:D, bottom).

Within the brown4 module, *Meg3* is the hub of a small gene network containing 16 additional long noncoding RNA (lncRNA) and 6 protein coding genes (Figure 7A). Two of the brown4 module lncRNA genes, *Meg3* and *Malat1* (blue*), were recently identified as direct targets of *Snord116* RNA-RNA interactions in mouse brain (Liu et al., 2024) (sex unspecified), an overlap that was significant (adjusted p=0.0235, Data S13). Five of the brown4 lncRNAs are parentally imprinted (Higgs et al., 2022), including three maternally expressed (Figure 7A pink*: *Meg3*, *B830012L14Rik*, *Ftx*) and two paternally expressed (*Kcnqlot1*, *A330076H08Rik*) genes, also significantly greater than expected (adjusted p=0.0045, Data S14). *A330076H08Rik* is a transcript extension of Zpf127 within the *Snord116* locus and *B830012L14Rik* localizes within the *Meg3* locus, as does *GM37899* from the brown4 module. *Ftx* is adjacent to *Xist* on the X chromosome and acts as an *Xist* enhancer (Chureau et al., 2011), but is dispensable for X chromosome inactivation (Soma et al., 2014). The *Snord116* host gene *116HG*, *Meg3*, *Xist*, and *Kcnq1ot1* are all imprinted in mouse and are characterized by chromatin-bound lncRNA clouds that bind to their site of transcription, but on four different imprinted loci (Clemson et al., 1996; Pandey et al., 2008; Powell et al., 2013a; Powell et al., 2013b; Sanli et al., 2018). A recent study using proximity labelling identified 98 RNA transcripts proximal to *Xist*, including 6 out of 22 transcripts from the brown4 module (Figure 7A green*: *Leng8, Spaca6, Ftx, Neat1, Malat1, Kcnq1ot1)* (Tsue et al., 2024), an enrichment that was highly significant (adjusted p value<0.0001, Data S15). Therefore, the brown4 correlated gene network is significantly enriched for sex-specific and imprinted lncRNAs with chromatin-bound nuclear organization and direct binding to *Snord116* RNA or *Xist* RNA.

**Figure 7.**
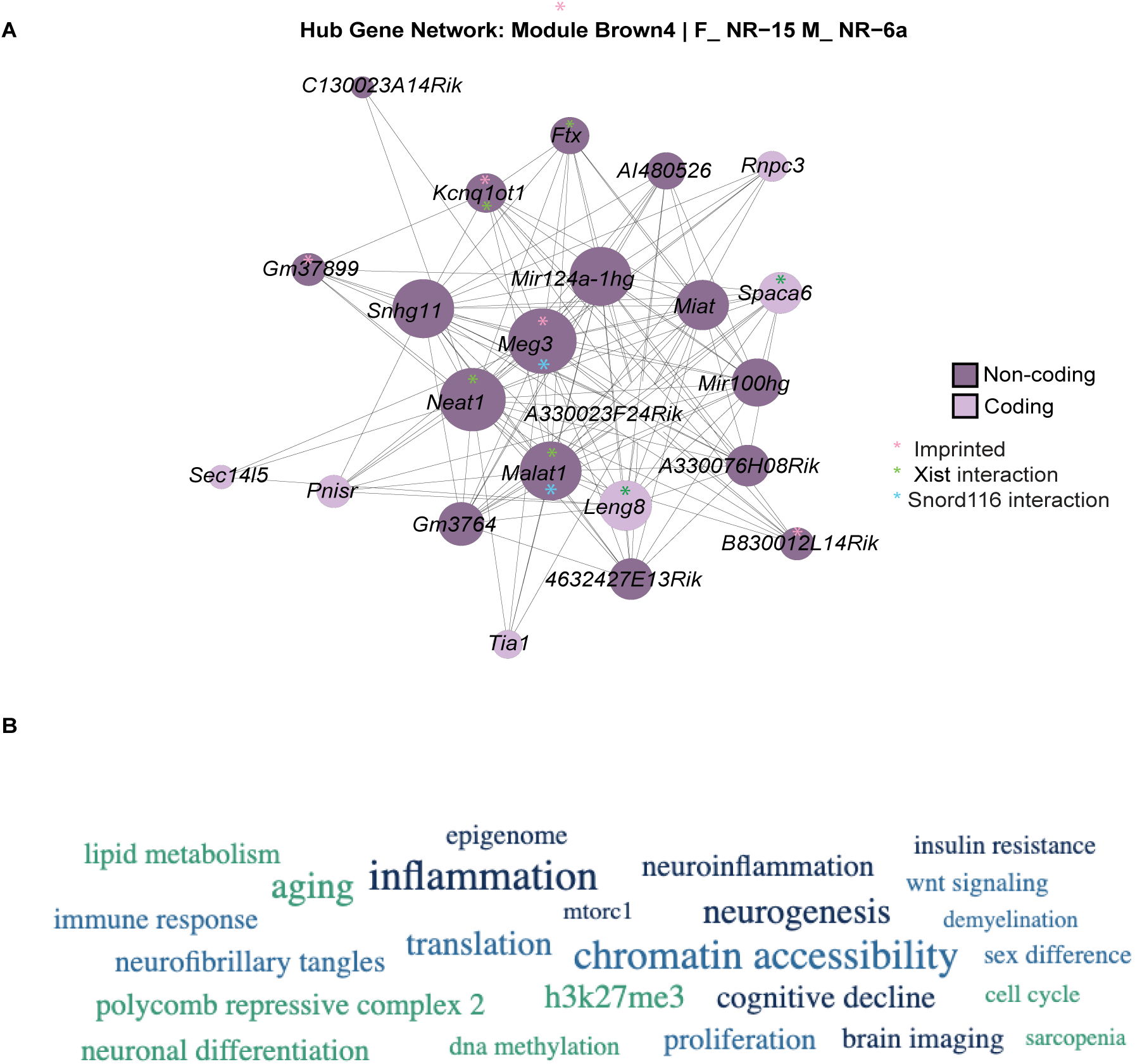
Sex- and light-sensitive module brown4, positively correlated with *Xist* expression and *Snord116* deletion in female mice, contains a small network of imprinted lncRNAs enriched for relevant biological pathways. (A) Module brown4 hub network shows *Meg3* as the top hub gene. LncRNAs, imprinted transcripts, and *Xist*-proximal transcripts are indicated. (B) The brown4 gene set was analyzed in RummaGEO for Pathway/Biological Process enriched terms, with the top 23 shown as a word cloud. Letter size reflects number of overlapping genes and darker colors represent greater significance. The top 50 terms and statistics are in Data S16.

The 22 genes in the brown4 module were analyzed for functional enrichments by RummaGEO Pathway/Biological Process terms, resulting in enrichment for terms relevant to our phenotypic results such as sex difference, circadian clock, lipid metabolism, and mTORC1, as well as epigenetic terms of chromatin accessibility, H3K27me3, and DNA methylation (Figure 7B, Data S16). Together, these results suggest that *Xist/Ftx* may modify the transcriptional and phenotypic effects of the *Snord116* deletion and transgene through a correlated network of imprinted regulatory lncRNAs implicated in neurologically relevant functions.

## Discussion

In this multifaceted investigation of genotype-phenotype correlations of the imprinted *Snord116* locus under dynamic diurnal lighting conditions, we have multiple new findings relevant to understanding factors influencing adaptation to a novel environment. First, we demonstrate that the effects of *Snord116* on transcriptional, metabolic, and behavioral phenotypes are modified by sex, light, and entrainment. Second, we demonstrate that a *Snord116* transgene expressed and localized outside of the endogenous imprinted locus can rescue a subset of the transcriptional changes of *Snord116* deficiency, but not the metabolic or behavioral phenotypes measured. Third, we created a rhythmic cortical transcriptional atlas of correlated gene networks that was used to compare gene networks across sexes and lighting conditions. Both *Snord116* deficiency and overexpression resulted in transcriptional changes in cortex to both rhythmic and non-rhythmic gene networks, with the largest genotype effects in diurnal L:D conditions but the largest entrainment effects in female cortex collected under constant darkness (D:D). Lastly, we identified a *Meg3*- and *Xist*-correlated gene network enriched for imprinted lncRNAs that may explain how the imprinted *Snord116* RNAs interact with female sex in the adaptation to a dynamic environment. These results are expected to be important in understanding the complex molecular pathogenesis of PWS as well as the function of *Snord116* more broadly in metabolic and neurologic diseases.

A major finding in our study was that genotype-phenotype correlations for *Snord116* were strongly influenced by extrinsic factors, including sex and light, in addition to T-cycle entrainment. While sex differences have been frequently observed in studies of diurnal rhythmicity of behavior and metabolism (Spitschan et al., 2022), they are not often investigated in PWS or any imprinted disease model. The female-specific effects of *Snord116* deletion mice showing lower overall activity and amplitude are consistent with other findings that describe wild-type female mice as showing higher overall free running activity compared to males (Lightfoot et al., 2004; Perrigo and Bronson, 1985) which has been attributed to estrogen (Krizo and Mintz, 2015; Lightfoot, 2008). However, estrogen levels are low in PWS females (Siemensma et al., 2012) and estrogen-related gene pathways were not particularly enriched in our analyses, suggesting a potential genetic X chromosomal contribution. Other studies have also indicated that female mice are more sensitive and adaptable to environmental cues and change their behavior in response to changes in diet and lighting conditions (Casimiro et al., 2021; Mei et al., 2021; Oraha et al., 2022; Smarr and Kriegsfeld, 2022). We have also previously demonstrated that cortical transcriptional and DNA methylation differences in *Snord116* deletion male mice were most apparent during the light hours (Coulson et al., 2018b; Powell et al., 2013b). The results of the current study are novel in demonstrating that the requirement for endogenous *Snord116* in behavioral adaptations to circadian entrainment differs by sex. However, our results are consistent with two studies of PWS in humans showing that female PWS patients are significantly more impacted in their metabolic measurements, while males have more profound behavioral deficits than females with PWS (Gito et al., 2015; Irizarry et al., 2015). Our results are also consistent with a study of healthy newborns showing that *MAGEL2, NDN*, *SNORD116*, and *SNORD115* expression in the umbilical cord was negatively associated with birth weight, length, and placental weight, with strongest effects in females (Mas-Pares et al., 2025). These combined results have implications for clinical considerations of potential sex differences in the treatment of PWS.

A major challenge for gene therapy for PWS is that the processing and localization of the *Snord116* snoRNAs to the nucleolus and the spliced host gene lncRNA to the site of transcription only occur at the endogenous locus, not from an exogenous locus (Coulson et al., 2018a). The current results of extensive phenotyping of PWS (HET/WT) and compensation (HET/TG) mice of both sexes were consistent with our prior study of body weight in male PWS and compensation mice in showing the lack of phenotypic rescue. However, we did observe a partial rescue of transcriptional changes associated with *Snord116* deletion in both sexes.

Furthermore, the *Snord116* transgene expressed on a wild-type background showed many significant phenotypic and transcriptional effects. These results can be explained by our prior fluorescence in situ hybridization results showing that the *Snord116* transgene was able to localize correctly only on the wild-type background, with the *116HG/Snhg14* lncRNA localizing to the endogenous locus RNA cloud (Coulson et al., 2018a). In the absence of the endogenous paternal *Snord116* locus, however, the transgenic transcript did not form a visible nuclear RNA cloud. The current results showing a partial transcriptional rescue in HET/TG mice suggests that some transcriptional roles of *Snord116* do not require localization to the active endogenous allele. However, future gene therapy strategies may need to consider the inherent molecular complexities of the PWS locus. Epigenetic-based therapies to activate the normally condensed chromatin state of the maternal *Snord116* locus may be one potential solution.

Despite the large energy burden of cellular metabolism within the cerebral cortex, this tissue has not been previously studied in circadian or diurnal transcriptomic investigations. Our study contributes a novel diurnal RNA-seq atlas of transcriptional regulation in the cerebral cortex through a 24 hour 12:12 L:D cycle for both sexes. We reduced the complexity of the diurnal cortical transcriptome by identifying rhythmic and non-rhythmic modules that were further used to examine transcriptional sex differences, as well as *Snord116* genotype-phenotype correlations across different entrainment and lighting conditions. Interestingly, gene modules that differed in rhythmicity between the sexes were significantly enriched for traits and diseases with described sex differences, results which are expected to be relevant to understanding sex differences in disease susceptibilities beyond PWS. In our PWS mouse model of paternal *Snord116* deficiency, transcriptional changes in cortex were observed in both rhythmic and non-rhythmic gene networks. Like what was observed with the behavioral and metabolic phenotypes, female cortex was transcriptionally more impacted by *Snord116* genotype changes than male cortex. Genotype effects were predominant in diurnal L:D conditions of light exposure in both males and females. In contrast, in conditions of constant darkness (D:D), females specifically were characterized by effects of both genotype as well as prior entrainment on transcriptional patterns that additionally correlated with metabolic phenotypes. These defined gene networks are an important first step in understanding how sex and light modify gene expression patterns in the cerebral cortex and more specifically in PWS.

Of the correlated gene networks we identified as being modified by *Snord116* genotype, compensation, sex differences, and/or lighting conditions, the brown4 module containing 22 coexpressed transcripts holds mechanistic insights into the complex relationships between genetic and non-genetic modifiers observed in this study. The imprinted lncRNA *Meg3* was the hub gene of the brown4 module and previously demonstrated to be altered in this PWS mouse model (Coulson et al., 2018b). *Meg3* resides within the *Dlk1/Dio3* imprinted locus that bears striking similarity to the PWS locus, as it encodes the only other repetitive cluster of snoRNAs in the mammalian genome (*SNORD113*, *SNORD114*), which are maternally expressed and exhibit allele-specific chromatin decondensation in neurons, similar to *SNORD116* and *SNORD115* (Cavaille et al., 2002; Leung et al., 2009; Tierling et al., 2006). Interestingly, loss of maternal *MEG3* in Temple syndrome phenocopies PWS, previously suggesting that these two imprinted loci may perform integrated functions and share common pathways (Hosoki et al., 2009; Kagami et al., 2015; Temple et al., 1991), including circadian (Devos et al., 2011; Kozlov et al., 2007; Labialle et al., 2008a; Labialle et al., 2008b; Tennese and Wevrick, 2011).

*MEG3* is highly expressed in cortex and implicated in neuronal MTORC1 signaling, apoptosis, metabolic processes, and long-term potentiation (Hamilton et al., 2020; Royer et al., 2022; Tan et al., 2017). *MEG3* overexpression was sufficient to induce necroptosis in a female human neuronal xenograft model of Alzheimer disease (Balusu et al., 2023). Cross-regulation between both imprinted loci *Snord116* and *Meg3* have been described previously (Coulson et al., 2018b; Liu et al., 2024; Stelzer et al., 2014), but mechanisms were previously poorly understood. Here, we demonstrate a novel correlated network of genes (brown4) connecting *Snord116* and *Meg3* with other imprinted genes *B830012L14Rik*, *Ftx*, *Kcnqlot1*, and *A330076H08Rik*. We further demonstrate that both sex and light modify the effect of *Snord116* on the *Meg3*-brown4 gene network in cortex. The strong correlation of the female-specific *Xist* transcript with brown4 expression, combined with the prior proximity mapping of *Xist* with 6 out of 22 of the brown4 transcripts (Tsue et al., 2024), strongly suggests that *Xist* may explain some of the sex differences observed in the transcriptional and metabolic phenotypes of *Snord116* deficiency and overexpression. These results also fit into emerging themes in the literature that *Xist* has nonconical functions beyond X chromosome inactivation (Dror et al., 2024; San Roman et al., 2023) and that imprinted genes can be co-regulated in networks (Baptissart et al., 2022; Lopez et al., 2017; Scagliotti et al., 2023). Future studies to determine more precise mechanisms of interactions between imprinted genes and *Xist* and their relevance to human metabolic traits and neurologic diseases are needed.

In conclusion, we demonstrate that genotype-phenotype correlations of the *Snord116* deletion mouse model of PWS is modified by both sex and lighting conditions and we further identified cortical gene transcriptional networks that correlate with these relationships. These results are important for the design of future gene therapies for PWS and have broader implications for understanding the mechanistic basis of sex differences in metabolic, sleep, and neurodegenerative disorders in humans.

## Acknowledgments

This work was supported by a National Institutes of Health (NIH) grant R01HD098038 to JML, the UC Davis Intellectual and Developmental Disabilities Research Center (IDDRC) P50HD103526, and the Mouse Metabolic Phenotyping Center (MMPC) Live at UC Davis (MMPCLive, RRID:SCR_015357, mmpc.ucdavis.edu) under the MMPCLive Program, grant U2CDK135074. The library preparation and sequencing was carried out by the DNA Technologies and Expression Analysis Cores at the UC Davis Genome Center and was supported by a NIH Shared Instrumentation Grant S10OD010786. We also thank Dr. Blythe Durbin-Johnson for statistical support and Dr. Joanna Chiu for circadian biology expertise.

## Author’s Contributions

JML, DHY, and JJR designed the study. JML acquired funding for the study and supervised the project. AJPM, JMR, KN, SH, OS, CD, CT, and DHY performed the mouse and laboratory work. JMR performed mouse phenotyping data analyses and statistics. AJM performed the bioinformatic analyses with help from CH. AJM and JML interpreted the results and wrote the manuscript. All authors reviewed, edited, and approved the final manuscript.

## STAR Methods

### Resource Availability

#### Lead Contact

Further information and requests for resources and reagents should be directed to and will be fulfilled by the lead contact, Janine M. LaSalle.

#### Materials Availability

This study did not generate new unique reagents.

### Data and Code Availability

- Raw and processed sequencing data has been deposited at GEO and is publicly available as of the date of publication. The Accession number is listed in the key resources table.
- All original code has been deposited at GitHub and is publicly available as of the date of publication. The URL and DOI are listed in the key resources table.
- Any additional information required to reanalyze the data reported in this paper is available from the lead contact upon request.

### Experimental Model and Subject Details

#### Mouse models, breeding, and housing conditions

Prader-Willi syndrome model *Snord116* deletion mice on a C57BL6/J background, described previously (Ding et al., 2005), were maintained in a breeding colony as heterozygous, maternally inherited mutants (*Snord116*^-/+^), which are asymptomatic because of exclusive paternal expression. PWS model mice of this line inherit a paternal deletion (*Snord116*^+/-^), which are referred to as HET in the Results section. A non-imprinted *Snord116* overexpression model (*Snord116 Ctg*) ubiquitously expresses a complete transgene containing 27 copies of *Snord116* exon-snoRNA-intron repeating units and has also been previously described (Coulson et al., 2018a). Both models were maintained as heterozygotes on a C57BL6J background under standard housing and 12 hour cycles of light and dark (12:12). For the experiments in this study, male maternally deleted *Snord116*^-/+^ mice were bred to female *Ctg*^+/-^ dams, so that litters contained equal mixtures of four different genotypes: wild-type for both: *Snord116*^+/+^;*Ctg*^-/-^ (WT/WT); PWS model: *Snord116*^+/-^;*Ctg*^-/-^ (Deletion, HET/WT); WT *Snord116*^+/+^;*Ctg*^+/-^ (Overexpression, WT/TG); and PWS *Snord116*^+/-^;*Ctg*^+/-^ (Compensation, HET/TG). All breeding and early post-weaning offspring were housed in 12:12 lighting conditions and post-weaning offspring were co-housed by sex. Genotypes were determined from tail snip DNA.

### Methods Details

#### Entrainment

Male and female offspring (WT/WT, *Snord116*^+/+^*Ctg*^-/-^; HET/WT, *Snord116*^+/-^*Ctg*^-/-^; WT/TG, *Snord116*^+/+^*Ctg*^+/-^; HET/TG, *Snord116*^+/-^*Ctg*^+/-^ were moved at ∼8 weeks of age into light-controlled incubator housing for a 7-day acclimation to the new housing. All mice were randomly divided into two entrainment groups, 12:12 or 11:11 light/dark (L:D) where they were housed in both entrainment groups at 22°C. After 15 days, one group of entrained mice were transferred to new cages equipped with running wheel actograms, where they remained at the entrained light cycles for an additional 7 days (22 d entrainment total). A different group of entrained mice were moved to indirect calorimetry chambers after 22 days of entrainment.

#### Running wheel behavioral activity measurements

After 15 days of entrainment, mice were transferred to standard shoebox cages with attached running wheels (Starr Life Sciences, Oakmont, PA) and assess designated light cycle parameters (11:11 or 12:12 light/dark). Voluntary running was measured for 7 days at 15 s intervals for 1 week (baseline) after which animals entrained to an 11:11 light cycle was transferred to 12:12 light cycle to assess running activity for an additional 7 days. Overlapping “first 4” and “last 4” day periods to examine after-effects. Running activity was measured as total, light and dark time and distance run. Individual running records were analyzed by cosinor rhythm analysis software (https://www.circadian.org/softwar.html, (Refinetti et al., 2007)) for three rhythmic parameters: mesor (mean), amplitude (half the range of daily excursion), and period. Following voluntary running assessment, mice were euthanized by exsanguination under isoflurane anesthesia under standard 12:12 light conditions and tissues collected at zeitgeiber time 6 (ZT6).

#### Indirect calorimetry

A subset of mice was entrained for an additional 7 days (22 days total) before assessing energy expenditure by indirect calorimetry for 7 days followed by body composition analysis. Energy expenditure by indirect calorimetry, ambulation (activity), and water and food intake were evaluated using an automated system (CLAMS: Comprehensive Lab Animal Monitoring System, Columbus Instruments, Columbus, OH). Mice were placed in acclimation chambers and housed in the entrainment environmental cabinet at 22°C for 24 hr. They were then transferred to calorimetry chambers inside a 22°C environmental cabinet and allowed to acclimate for 24 hours prior to the start of the calorimetry measurements. Both acclimation (24 h) and calorimetry measurements (48 h) were collected at the entrained light cycles (11:11 or 12:12). After this 48-hour period, the light cycle was switched to total (0:24) darkness (D:D) and calorimetry measurements were collected for an additional 120 hours (4 days). Energy expenditure (kcal) was calculated from O_2_ consumption and CO_2_ production using the CLAMS Oxymax software. Respiratory exchange ratio (RER) was calculated as VCO_2_/VO_2_. Room air was drawn through the system at a rate of 500 ml/min. The oxygen and carbon dioxide analyzers were calibrated daily. A calibration gas (0.50% CO_2_, 20.50% O_2_, and balance nitrogen) (Airgas, Sacramento, CA) and dry room air were used to calibrate the analyzers. At the beginning and end of the experiment, the performance of the entire calorimetry system was assessed by bleeding a 20% CO_2_ (balance nitrogen) standard (Airgas, Sacramento, CA) through each chamber at a regulated rate using an OxyVal gas infusion system (Columbus Instruments, Columbus, OH) and measuring recovery of CO_2_ and dilution of O_2_ in the exhaust flow from the chambers. A cage-mounted infrared photocell system (Columbus Instruments, Columbus, OH) was used to assess activity (X & Z beam breaks). At the conclusion of the indirect calorimetry measurements, body composition (lean mass, fat mass, bone mineral content and bone mineral density) was assessed under isoflurane (2-5%) anesthesia and red light conditions using a Lunar PIXImus II Densitometer, (GE Medical Systems, Chalfont St. Giles, UK). Mice were then immediately euthanized by exsanguination under isoflurane anesthesia under red light and tissues collected at zeitgeiber time 6 (ZT6).

#### RNA Sequencing

Wild-type mice housed in home cage 12:12 light cycle were sacrificed every three hours at zeitgeiber time 0 – 21 (ZT0, ZT3, ZT6, ZT9, ZT12, ZT15, ZT18, ZT21) to generate a circadian atlas for the prefrontal cortex. RNA were isolated from 10 - 20 mg prefrontal cortex using All Prep DNA/RNA/Protein kit (Qiagen). RNA was submitted for library preparation and sequencing to Novogene. Libraries were prepared using NEBNext Ultra II Directional RNA library prep kit for Illumina. Ribosomal RNA was cleaned up using NEBNext rRNA Depletion kit. Libraries were sequenced on the Novaseq S4. RNA was also collected from ZT6 prefrontal cortex from indirect calorimetry and voluntary running experiments and submitted to Novogene for library prepration and sequencing. On average 20 million reads were produced for each sequencing experiment. Reads were processed using TrimGalore, Samtools, and STAR (https://github.com/ben-laufer/RNA-seq). Raw counts from each experiment were experiment and sex stratified and normalized (logcpm) using Limma-Voom. After normalization, sequencing samples were merged using R’s merge function to generate a large data frame that preserves all genes expressed across experiments. The final samples consisted of 6 groups: 32 D:D female and 31 D:D male (from indirect calorimetry experiment), 32 L:D female and 32 L:D male (from voluntary running experiment), and 46 wild-type female and 46 wild-type male (L:D 12:12 home cage, circadian atlas experiment) samples totaling 15,551 genes that were used for universal gene network analyses.

### Quantification and Statistical Analyses

#### Statistical Analyses of Phenotypes

Statistical analysis of measurements from indirect calorimetry and voluntary running were analyzed using two- and three-way ANOVA within PRISM. The interaction between genotype, sex, and entrainment were tested for both voluntary running and indirect calorimetry. In addition, the interaction between genotype, sex, and light treatment was also tested for indirect calorimetry since measurements were collected under constant dark conditions (D:D). Two-way ANOVA statistics were sex-stratified to identify genotype specific effects along with after-effects of treatments based on shifts of entrainment or lighting conditions. Multiple comparisons were performed to identify significant genotype differences. Because of the expected strong correlations between phenotypic measurements, they were not considered independent hypotheses, so no adjustments for multiple comparisons were performed.

#### Weighted Gene Correlation Network Analysis (WGCNA)

Normalized RNAseq counts were analyzed using consensus WGCNA R package (Langfelder and Horvath, 2008) to identify shared and unique networks across each sequencing experiment. Consensus modules were generated using the WGCNA settings adapted from (Mordaunt et al., 2019) using a signed network and a soft power threshold of 16. Expression data was checked for any missing data and imputed to preserve the uniquely expressed genes in each data set. Of the 15,551 genes, 2062 genes were excluded from having too many missing values. These genes were missing in the universal gene set because of their uniqueness to each sample set (such as *Xist* expression in females). Signed topological overlap matrices (TOMs) were generated for each individual experiment for the remaining 13,489 genes. TOMs were calibrated using a full quantile normalization to generate a consensus TOM in a single block representing overlaps across each study. String networks were generated using TOM matrices in cytoscape. None of the modules were merged to preserve the complexity produced by the circadian atlas data sets. The top 5 hub genes were determined in each module for each experiment by filtering the top 5 genes with the highest module membership. The measurements collected from CLAMS (indirect calorimetry) and the running wheel (voluntary running) experiments were used as traits correlated with the module eigengenes. Individual module trait correlations were produced using spearman’s method and pairwise complete observations. Fisher’s asymptotic was used to acquire p-values using a two-sided test. Significant correlations were determined with a p-value < 0.05. Aside from module-trait correlations, module-module, and trait-trait correlations were carried out to determine module-trait-genotype relationships. Module-module and trait-trait correlations were also carried out using Spearman’s method and pairwise complete observations. P-values were acquired using Fisher’s asymptotic using a two-sided test. Because of the high level of observed and expected module-module and trait-trait correlations, we did not adjust for multiple hypotheses and instead consider the strength of each correlation in the interpretation.

#### Circadian Statistical Analysis

Circadian analysis was performed using RAIN circadian statistics (Thaben and Westermark, 2014) to determine module eigenvalue rhythmicity of wild-type circadian atlas RNAseq datasets. RAIN analysis was sex stratified using options deltat = 3, period = 24, measure.sequence = c(6, 6, 6, 6, 6, 6, 5, 5), peak.border = c(0.3, 0.7), verbose = FALSE. Output of the analysis was used to rename modules and create heatmaps that show module eigenvalue rhythms. The module order of both heatmaps follow the rhythmicity of female modules.

#### Gene Function and Human Phenotype Enrichment Analyses

KEGG, dbGaP, and GO analysis was performed using EnrichR3.2 (Chen et al., 2013) for genes in each module. Terms are separated by rhythmic and non-rhythmic modules determined by the circadian analysis. KEGG and dbGaP dotplots are generated using ggplot and only shows terms with an adjusted p-value < 0.05. Supplementary tables of KEGG, dbGaP, and GO terms only show terms with an adjusted p-value < 0.05. Small gene sets were insufficient for KEGG, dbGaP, and GO analysis were analyzed using RummaGEO (Marino et al., 2024). RummaGEO figures of disease & phenotypes and pathways & biological processes only show terms with an adjusted p-value < 0.05.

#### Network Analysis and Visualization

Input data was acquired from conensusWGCNA consensus TOM outputs and exported to Cytoscape for gene network visualization (Shannon et al., 2003). Module-trait Sankey networks are generated using correlation coefficients and p-values. Module-trait networks were visualized using ggalluvial to show the frequency and distribution of significant correlations.

## Supplemental Information Titles and Legends Supplementary Figure file, containing S1-S6

**Figure S1. Female-specific effects on total running duration and distance.** (**A**) Scatter plot of total running time measurements for each genotype and sex taken at week 1 and week 2 with 3-way ANOVA statistics of entrainment, genotype, and sex. (**B**) Scatter plot of total running distance measurements for each genotype and sex taken at week 1 and week 2 with 3-way ANOVA statistics of entrainment, genotype, and sex. Open symbols, 11:11; closed symbols, 12:12; blue symbols, male; purple symbols, female.

**Figure S2. *Snord116* transgene does not rescue the reduced body weight phenotype in *Snord116* deletion mice.** (**A**) Body weight (BW). (B) % fat. (C) Fat mass. (D) Lean mass measurements taken for each genotype and sex separated by entrainment 11 (white) and 12 (skyblue – males, pink – females). (E) 2-way ANOVA statistics table of genotype, entrainment, and interaction effects of DEXA measurements including bone mineral density (BMD) and bone mineral composition (BMC).

**Figure S3. Sex-specific metabolic adaptations to entrainment and *Snord116* genotype.**

**(A)** L:D and (**B**) D:D scatter plots of energy expenditure (heat) measurements for each genotype and sex taken during light hours and dark hours with 3-way ANOVA statistics of entrainment, genotype, and sex. (**C**) L:D and (**D**) D:D scatter plots of food intake (kcal/hr) measurements for each genotype and sex taken during light hours and dark hours with 3-way ANOVA statistics of entrainment, genotype, and sex. (**E**) L:D and (**F**) D:D scatter plots of inactivity (% sleep) measurements for each genotype and sex taken during light hours and dark hours with 3-way ANOVA statistics of entrainment, genotype, and sex.

**Figure S4. Sex-specific metabolic adaptations to variable lighting conditions and *Snord116* genotype.** (**A**) 12 and (**B**) 11 scatter plots of respiratory exchange rate (RER) measurements for each genotype and sex taken during light hours and dark hours with 3-way ANOVA statistics of light treatment, genotype, and sex. (**C**) 12 and (**D**) 11 scatter plots of energy expenditure (heat) measurements for each genotype and sex taken during light hours and dark hours with 3-way ANOVA statistics of light treatment, genotype, and sex. (**E**) 12 and (**F**) 11 scatter plots of energy intake (kcal/hr) measurements for each genotype and sex taken during light hours and dark hours with 3-way ANOVA statistics of light treatment, genotype, and sex. (**G**) 12 and (**H**) 11 scatter plots of activity (X-tot) measurements for each genotype and sex taken during light hours and dark hours with 3-way ANOVA statistics of light treatment, genotype, and sex. (**I**) 12 and (**J**) 11 scatter plots of inactivity (% sleep) measurements for each genotype and sex taken during light hours and dark hours with 3-way ANOVA statistics of light treatment, genotype, and sex.

**Figure S5. WGCNA module networks are highly preserved across experiments.** Module preservation statistics of each experimental data set. Includes wild-type circadian atlas male and female, CLAMS male and female, and running wheel male and female normalized (logcpm) RNAseq counts. The density value (*D*) indicates the preservation score across all datasets. Any value *D* > 0.5 is considered highly preserved.

**Figure S6. Module networks highly correlated with the *Snord116* transgene are enriched for terms relevant to PWS.** (**A**) Dotplot of database of genotype and phenotypes (dbGaP) terms for modules with significant (Adjusted.pvalue < 0.05) terms. (**B**) Dotplot of top 100 KEGG for modules with significant (Adjusted.pvalue < 0.05) terms.

## Supplementary Data file, containing Data S1-S16 in tabbed .xlxs file

**Data S1:** Numbers of mice of each genotype and sex obtained from breeding with statistics

**Data S2:** Running wheel phenotypes (all) statistical analyses with p values from pairwise comparisons of each genotype within each sex and entrainment group

**Data S3:** Running wheel phenotypes (light versus dark hours) statistical analyses with p values from pairwise comparisons of each genotype within each sex and entrainment group

**Data S4:** CLAMS phenotypes (light versus dark hours) statistical analyses with p values from pairwise comparisons of each genotype within each sex and entrainment group

**Data S5:** Statistical analyses with p values for differences in body composition scores by genotype and entrainment, sex-stratified

**Data S6:** Top 5 hub genes for each WGCNA module defined in Figures 4-5 by sex and lighting condition

**Data S7:** Significantly enriched GO terms for Biological Process for each WGCNA module

**Data S8:** Significantly enriched GO terms for Cellular Component for each WGCNA module

**Data S9:** Significantly enriched GO terms for Molecular Function for each WGCNA module

**Data S10:** Significantly enriched KEGG Pathways for each WGCNA module

**Data S11:** Significantly enriched dbGAP terms for each WGCNA module

**Data S12:** Significantly enriched RNADD GEO experiments for each WGCNA module

**Data S13:** Module overlap with lint of imprinted genes in mouse brain from PMID: 36384442

**Data S14:** Module overlap with Xist interacting RNA from O-MAP-Seq in PMID: 39468212

**Data S15:** *Snord116* bound RNAs from mouse brain (sex not specified) in PMID: 39579764

**Data S16:** brown4 module enriched in RummaGEO Pathway/Biological Pathway terms from PMID: 39569206

## References

Balusu, S., Horre, K., Thrupp, N., Craessaerts, K., Snellinx, A., Serneels, L., T’Syen, D., Chrysidou, I., Arranz, A.M., Sierksma, A., et al. (2023). MEG3 activates necroptosis in human neuron xenografts modeling Alzheimer’s disease. Science 381, 1176–1182.

Baptissart, M., Bradish, C.M., Jones, B.S., Walsh, E., Tehrani, J., Marrero-Colon, V., Mehta, S., Jima, D.D., Oh, S.H., Diehl, A.M., et al. (2022). Zac1 and the Imprinted Gene Network program juvenile NAFLD in response to maternal metabolic syndrome. Hepatology 76, 1090–1104.

Bieth, E., Eddiry, S., Gaston, V., Lorenzini, F., Buffet, A., Conte Auriol, F., Molinas, C., Cailley, D., Rooryck, C., Arveiler, B., et al. (2015). Highly restricted deletion of the SNORD116 region is implicated in Prader-Willi Syndrome. European journal of human genetics : EJHG 23, 252–255.

Blasiak, A., Gundlach, A.L., Hess, G., and Lewandowski, M.H. (2017). Interactions of Circadian Rhythmicity, Stress and Orexigenic Neuropeptide Systems: Implications for Food Intake Control. Front Neurosci 11, 127.

Bronstein, P.M., Wolkoff, F.D., and Levine, M.J. (1975). Sex-related differences in rats’ open-field activity. Behavioral Biology 13, 133–138.

Burnett, L.C., Hubner, G., LeDuc, C.A., Morabito, M.V., Carli, J.F.M., and Leibel, R.L. (2017). Loss of the imprinted, non-coding Snord116 gene cluster in the interval deleted in the Prader Willi syndrome results in murine neuronal and endocrine pancreatic developmental phenotypes. Hum Mol Genet 26, 4606–4616.

Butler, J.V., Whittington, J.E., Holland, A.J., Boer, H., Clarke, D., and Webb, T. (2002). Prevalence of, and risk factors for, physical ill-health in people with Prader-Willi syndrome: a population-based study. Dev Med Child Neurol 44, 248–255.

Casimiro, I., Stull, N.D., Tersey, S.A., and Mirmira, R.G. (2021). Phenotypic sexual dimorphism in response to dietary fat manipulation in C57BL/6J mice. Journal of Diabetes and its Complications 35, 107795.

Cassidy, S.B., Schwartz, S., Miller, J.L., and Driscoll, D.J. (2012). Prader-Willi syndrome. Genetics in medicine : official journal of the American College of Medical Genetics 14, 10–26.

Cavaille, J., Seitz, H., Paulsen, M., Ferguson-Smith, A.C., and Bachellerie, J.P. (2002). Identification of tandemly-repeated C/D snoRNA genes at the imprinted human 14q32 domain reminiscent of those at the Prader-Willi/Angelman syndrome region. Hum Mol Genet 11, 1527–1538.

Chen, E.Y., Tan, C.M., Kou, Y., Duan, Q., Wang, Z., Meirelles, G.V., Clark, N.R., and Ma’ayan, A. (2013). Enrichr: interactive and collaborative HTML5 gene list enrichment analysis tool. BMC Bioinformatics 14, 128.

Chureau, C., Chantalat, S., Romito, A., Galvani, A., Duret, L., Avner, P., and Rougeulle, C. (2011). Ftx is a non-coding RNA which affects Xist expression and chromatin structure within the X-inactivation center region. Hum Mol Genet 20, 705–718.

Clemson, C.M., McNeil, J.A., Willard, H.F., and Lawrence, J.B. (1996). XIST RNA paints the inactive X chromosome at interphase: evidence for a novel RNA involved in nuclear/chromosome structure. J Cell Biol 132, 259–275.

Colas, D., Wagstaff, J., Fort, P., Salvert, D., and Sarda, N. (2005). Sleep disturbances in Ube3a maternal-deficient mice modeling Angelman syndrome. Neurobiology of disease 20, 471–478.

Coulson, R.L., Powell, W.T., Yasui, D.H., Dileep, G., Resnick, J., and LaSalle, J.M. (2018a). Prader-Willi locus Snord116 RNA processing requires an active endogenous allele and neuron-specific splicing by Rbfox3/NeuN. Human molecular genetics 27, 4051–4060.

Coulson, R.L., Yasui, D.H., Dunaway, K.W., Laufer, B.I., Vogel Ciernia, A., Zhu, Y., Mordaunt, C.E., Totah, T.S., and Lasalle, J.M. (2018b). Snord116-dependent diurnal rhythm of DNA methylation in mouse cortex. Nat Commun 9, 1616.

de Smith, A.J., Purmann, C., Walters, R.G., Ellis, R.J., Holder, S.E., Van Haelst, M.M., Brady, A.F., Fairbrother, U.L., Dattani, M., Keogh, J.M., et al. (2009). A Deletion of the HBII-85 Class of Small Nucleolar RNAs (snoRNAs) is Associated with Hyperphagia, Obesity and Hypogonadism. Hum Mol Genet 18, 3257–3265.

Devos, J., Weselake, S.V., and Wevrick, R. (2011). Magel2, a Prader-Willi syndrome candidate gene, modulates the activities of circadian rhythm proteins in cultured cells. J Circadian Rhythms 9, 12.

Ding, F., Li, H.H., Zhang, S., Solomon, N.M., Camper, S.A., Cohen, P., and Francke, U. (2008). SnoRNA Snord116 (Pwcr1/MBII-85) deletion causes growth deficiency and hyperphagia in mice. PLoS ONE 3, e1709.

Ding, F., Prints, Y., Dhar, M.S., Johnson, D.K., Garnacho-Montero, C., Nicholls, R.D., and Francke, U. (2005). Lack of Pwcr1/MBII-85 snoRNA is critical for neonatal lethality in Prader-Willi syndrome mouse models. Mamm Genome 16, 424–431.

Dror, I., Chitiashvili, T., Tan, S.Y.X., Cano, C.T., Sahakyan, A., Markaki, Y., Chronis, C., Collier, A.J., Deng, W., Liang, G., et al. (2024). XIST directly regulates X-linked and autosomal genes in naive human pluripotent cells. Cell 187, 110–129 e131.

Duker, A.L., Ballif, B.C., Bawle, E.V., Person, R.E., Mahadevan, S., Alliman, S., Thompson, R., Traylor, R., Bejjani, B.A., Shaffer, L.G., et al. (2010). Paternally inherited microdeletion at 15q11.2 confirms a significant role for the SNORD116 C/D box snoRNA cluster in Prader-Willi syndrome. Eur J Hum Genet 18, 1196–1201.

Esbensen, A.J., and Schwichtenberg, A.J. (2016). Sleep in Neurodevelopmental Disorders. Int Rev Res Dev Disabil 51, 153–191.

Fischer, K.E., Hoffman, J.M., Sloane, L.B., Gelfond, J.A.L., Soto, V.Y., Richardson, A.G., and Austad, S.N. (2016). A cross-sectional study of male and female C57BL/6Nia mice suggests lifespan and healthspan are not necessarily correlated. Aging 8, 2370–2391.

Gibbs, S., Wiltshire, E., and Elder, D. (2013). Nocturnal sleep measured by actigraphy in children with Prader-Willi syndrome. The Journal of pediatrics 162, 765–769.

Gito, M., Ihara, H., Ogata, H., Sayama, M., Murakami, N., Nagai, T., Ayabe, T., Oto, Y., and Shimoda, K. (2015). Gender Differences in the Behavioral Symptom Severity of Prader-Willi Syndrome. Behav Neurol 2015, 294127.

Hamilton, S., de Cabo, R., and Bernier, M. (2020). Maternally expressed gene 3 in metabolic programming. Biochim Biophys Acta Gene Regul Mech 1863, 194396.

Herculano-Houzel, S. (2011). Scaling of brain metabolism with a fixed energy budget per neuron: implications for neuronal activity, plasticity and evolution. PLoS One 6, e17514.

Higgs, M.J., Hill, M.J., John, R.M., and Isles, A.R. (2022). Systematic investigation of imprinted gene expression and enrichment in the mouse brain explored at single-cell resolution. BMC Genomics 23, 754.

Hosoki, K., Kagami, M., Tanaka, T., Kubota, M., Kurosawa, K., Kato, M., Uetake, K., Tohyama, J., Ogata, T., and Saitoh, S. (2009). Maternal uniparental disomy 14 syndrome demonstrates prader-willi syndrome-like phenotype. J Pediatr 155, 900–903 e901.

Irizarry, K.A., Bain, J., Butler, M.G., Ilkayeva, O., Muehlbauer, M., Haqq, A.M., and Freemark, M. (2015). Metabolic profiling in Prader-Willi syndrome and nonsyndromic obesity: sex differences and the role of growth hormone. Clin Endocrinol (Oxf) 83, 797–805.

Kagami, M., Kurosawa, K., Miyazaki, O., Ishino, F., Matsuoka, K., and Ogata, T. (2015). Comprehensive clinical studies in 34 patients with molecularly defined UPD(14)pat and related conditions (Kagami-Ogata syndrome). Eur J Hum Genet 23, 1488–1498.

Koike, N., Yoo, S.H., Huang, H.C., Kumar, V., Lee, C., Kim, T.K., and Takahashi, J.S. (2012). Transcriptional architecture and chromatin landscape of the core circadian clock in mammals. Science 338, 349–354.

Kozlov, S.V., Bogenpohl, J.W., Howell, M.P., Wevrick, R., Panda, S., Hogenesch, J.B., Muglia, L.J., Van Gelder, R.N., Herzog, E.D., and Stewart, C.L. (2007). The imprinted gene Magel2 regulates normal circadian output. Nat Genet 39, 1266–1272.

Krizo, J.A., and Mintz, E.M. (2015). Sex Differences in Behavioral Circadian Rhythms in Laboratory Rodents. Front Endocrinol 5.

Labialle, S., Croteau, S., Belanger, V., McMurray, E.N., Ruan, X., Moussette, S., Jonnaert, M., Schmidt, J.V., Cermakian, N., and Naumova, A.K. (2008a). Novel imprinted transcripts from the Dlk1-Gtl2 intergenic region, Mico1 and Mico1os, show circadian oscillations. Epigenetics 3, 322–329.

Labialle, S., Yang, L., Ruan, X., Villemain, A., Schmidt, J.V., Hernandez, A., Wiltshire, T., Cermakian, N., and Naumova, A.K. (2008b). Coordinated diurnal regulation of genes from the Dlk1-Dio3 imprinted domain: implications for regulation of clusters of non-paralogous genes. Human Molecular Genetics 17, 15–26.

Langfelder, P., and Horvath, S. (2008). WGCNA: an R package for weighted correlation network analysis. BMC Bioinformatics 9, 559.

Lassi, G., Priano, L., Maggi, S., Garcia-Garcia, C., Balzani, E., El-Assawy, N., Pagani, M., Tinarelli, F., Giardino, D., Mauro, A., et al. (2016). Deletion of the Snord116/SNORD116 Alters Sleep in Mice and Patients with Prader-Willi Syndrome. Sleep 39, 637–644.

Legates, T.A., Fernandez, D.C., and Hattar, S. (2014). Light as a central modulator of circadian rhythms, sleep and affect. Nature Reviews Neuroscience 15, 443–454.

Leung, K.N., Vallero, R.O., DuBose, A.J., Resnick, J.L., and LaSalle, J.M. (2009). Imprinting regulates mammalian snoRNA-encoding chromatin decondensation and neuronal nucleolar size. Hum Mol Genet 18, 4227–4238.

Lightfoot, J.T. (2008). Sex Hormones’ Regulation of Rodent Physical Activity: A Review. Int J Biol Sci, 126–132.

Lightfoot, J.T., Turner, M.J., Daves, M., Vordermark, A., and Kleeberger, S.R. (2004). Genetic influence on daily wheel running activity level. Physiol Genomics 19, 270–276.

Liu, B., Wu, T., Miao, B.A., Ji, F., Liu, S., Wang, P., Zhao, Y., Zhong, Y., Sundaram, A., Zeng, T.B., et al. (2024). snoRNA-facilitated protein secretion revealed by transcriptome-wide snoRNA target identification. Cell.

Lopez, S.J., Dunaway, K., Islam, M.S., Mordaunt, C., Vogel Ciernia, A., Meguro-Horike, M., Horike, S.I., Segal, D.J., and LaSalle, J.M. (2017). UBE3A-mediated regulation of imprinted genes and epigenome-wide marks in human neurons. Epigenetics 12, 982–990.

Marino, G.B., Clarke, D.J.B., Deng, E.Z., and Ma’ayan, A. (2024). RummaGEO: Automatic Mining of Human and Mouse Gene Sets from GEO. bioRxiv: The Preprint Server for Biology, 2024.2004.2009.588712.

Mas-Pares, B., Carreras-Badosa, G., Gomez-Vilarrubla, A., De Arriba-Munoz, A., Lafalla-Bernard, O., Prats-Puig, A., De Zegher, F., Ibanez, L., Haqq, A.M., Bassols, J., et al. (2025). Sex dimorphic associations of Prader-Willi imprinted gene expressions in umbilical cord with prenatal and postnatal growth in healthy infants. World J Pediatr 21, 100–112.

Mei, Y., Teng, H., Li, Z., Zeng, C., Li, Y., Song, W., Zhang, K., Sun, Z.S., and Wang, Y. (2021). Restricted Feeding Resets Endogenous Circadian Rhythm in Female Mice Under Constant Darkness. Neurosci Bull 37, 1005–1009.

Mordaunt, C.E., Park, B.Y., Bakulski, K.M., Feinberg, J.I., Croen, L.A., Ladd-Acosta, C., Newschaffer, C.J., Volk, H.E., Ozonoff, S., Hertz-Picciotto, I., et al. (2019). A meta-analysis of two high-risk prospective cohort studies reveals autism-specific transcriptional changes to chromatin, autoimmune, and environmental response genes in umbilical cord blood. Mol Autism 10, 36.

Mukherji, A., Kobiita, A., Damara, M., Misra, N., Meziane, H., Champy, M.F., and Chambon, P. (2015). Shifting eating to the circadian rest phase misaligns the peripheral clocks with the master SCN clock and leads to a metabolic syndrome. Proc Natl Acad Sci U S A 112, E6691–6698.

Oraha, J., Enriquez, R.F., Herzog, H., and Lee, N.J. (2022). Sex-specific changes in metabolism during the transition from chow to high-fat diet feeding are abolished in response to dieting in C57BL/6J mice. Int J Obes (Lond) 46, 1749–1758.

Pandey, R.R., Mondal, T., Mohammad, F., Enroth, S., Redrup, L., Komorowski, J., Nagano, T., Mancini-Dinardo, D., and Kanduri, C. (2008). Kcnq1ot1 antisense noncoding RNA mediates lineage-specific transcriptional silencing through chromatin-level regulation. Mol Cell 32, 232–246.

Papazyan, R., Zhang, Y., and Lazar, M.A. (2016). Genetic and epigenomic mechanisms of mammalian circadian transcription. Nat Struct Mol Biol 23, 1045–1052.

Perrigo, G., and Bronson, F.H. (1985). Sex differences in the energy allocation strategies of house mice. Behav Ecol Sociobiol 17, 297–302.

Powell, W.T., Coulson, R.L., Crary, F.K., Wong, S.S., Ach, R.A., Tsang, P., Yamada, N.A., Yasui, D.H., and LaSalle, J.M. (2013a). A Prader-Willi locus lncRNA cloud modulates diurnal genes and energy expenditure. Human Molecular Genetics 22, 4318–4328.

Powell, W.T., Coulson, R.L., Gonzales, M.L., Crary, F.K., Wong, S.S., Adams, S., Ach, R.A., Tsang, P., Yamada, N.A., Yasui, D.H., et al. (2013b). R-loop formation at Snord116 mediates topotecan inhibition of Ube3a-antisense and allele-specific chromatin decondensation. Proceedings of the National Academy of Sciences of the United States of America 110, 13938–13943.

Qian, P., He, X.C., Paulson, A., Li, Z., Tao, F., Perry, J.M., Guo, F., Zhao, M., Zhi, L., Venkatraman, A., et al. (2016). The Dlk1-Gtl2 Locus Preserves LT-HSC Function by Inhibiting the PI3K-mTOR Pathway to Restrict Mitochondrial Metabolism. Cell Stem Cell 18, 214–228.

Refinetti, R., Lissen, G.C., and Halberg, F. (2007). Procedures for numerical analysis of circadian rhythms. Biol Rhythm Res 38, 275–325.

Rosenfeld, C.S. (2017). Sex-dependent differences in voluntary physical activity. J of Neuroscience Research 95, 279–290.

Royer, M., Pai, B., Menon, R., Bludau, A., Gryksa, K., Perry, R.B., Ulitsky, I., Meister, G., and Neumann, I.D. (2022). Transcriptome and chromatin alterations in social fear indicate association of MEG3 with successful extinction of fear. Mol Psychiatry 27, 4064–4076.

Sahoo, T., del Gaudio, D., German, J.R., Shinawi, M., Peters, S.U., Person, R.E., Garnica, A., Cheung, S.W., and Beaudet, A.L. (2008). Prader-Willi phenotype caused by paternal deficiency for the HBII-85 C/D box small nucleolar RNA cluster. Nat Genet 40, 719–721.

San Roman, A.K., Godfrey, A.K., Skaletsky, H., Bellott, D.W., Groff, A.F., Harris, H.L., Blanton, L.V., Hughes, J.F., Brown, L., Phou, S., et al. (2023). The human inactive X chromosome modulates expression of the active X chromosome. Cell Genomics 3, 100259.

Sanli, I., Lalevee, S., Cammisa, M., Perrin, A., Rage, F., Lleres, D., Riccio, A., Bertrand, E., and Feil, R. (2018). Meg3 Non-coding RNA Expression Controls Imprinting by Preventing Transcriptional Upregulation in cis. Cell Rep 23, 337–348.

Scagliotti, V., Vignola, M.L., Willis, T., Howard, M., Marinelli, E., Gaston-Massuet, C., Andoniadou, C., and Charalambous, M. (2023). Imprinted Dlk1 dosage as a size determinant of the mammalian pituitary gland. Elife (Cambridge) 12.

Shannon, P., Markiel, A., Ozier, O., Baliga, N.S., Wang, J.T., Ramage, D., Amin, N., Schwikowski, B., and Ideker, T. (2003). Cytoscape: A Software Environment for Integrated Models of Biomolecular Interaction Networks. Genome Research 13, 2498–2504.

Siemensma, E.P., van Alfen-van der Velden, A.A., Otten, B.J., Laven, J.S., and Hokken-Koelega, A.C. (2012). Ovarian function and reproductive hormone levels in girls with Prader-Willi syndrome: a longitudinal study. J Clin Endocrinol Metab 97, E1766–1773.

Smarr, B., and Kriegsfeld, L.J. (2022). Female mice exhibit less overall variance, with a higher proportion of structured variance, than males at multiple timescales of continuous body temperature and locomotive activity records. Biology of Sex Differences 13.

Soma, M., Fujihara, Y., Okabe, M., Ishino, F., and Kobayashi, S. (2014). Ftx is dispensable for imprinted X-chromosome inactivation in preimplantation mouse embryos. Sci Rep 4, 5181.

Spitschan, M., Santhi, N., Ahluwalia, A., Fischer, D., Hunt, L., Karp, N.A., Levi, F., Pineda-Torra, I., Vidafar, P., and White, R. (2022). Sex differences and sex bias in human circadian and sleep physiology research. Elife (Cambridge) 11.

Stelzer, Y., Sagi, I., Yanuka, O., Eiges, R., and Benvenisty, N. (2014). The noncoding RNA IPW regulates the imprinted DLK1-DIO3 locus in an induced pluripotent stem cell model of Prader-Willi syndrome. Nature Genetics 46, 551–557.

Takahashi, J.S. (2017). Transcriptional architecture of the mammalian circadian clock. Nat Rev Genet 18, 164–179.

Tan, M.C., Widagdo, J., Chau, Y.Q., Zhu, T., Wong, J.J., Cheung, A., and Anggono, V. (2017). The Activity-Induced Long Non-Coding RNA Meg3 Modulates AMPA Receptor Surface Expression in Primary Cortical Neurons. Front Cell Neurosci 11, 124.

Temple, I.K., Cockwell, A., Hassold, T., Pettay, D., and Jacobs, P. (1991). Maternal uniparental disomy for chromosome 14. J Med Genet 28, 511–514.

Tennese, A.A., and Wevrick, R. (2011). Impaired hypothalamic regulation of endocrine function and delayed counterregulatory response to hypoglycemia in Magel2-null mice. Endocrinology 152, 967–978.

Thaben, P.F., and Westermark, P.O. (2014). Detecting rhythms in time series with rain. Journal of Biological Rhythms.

Thibert, R.L., Larson, A.M., Hsieh, D.T., Raby, A.R., and Thiele, E.A. (2013). Neurologic manifestations of Angelman syndrome. Pediatric neurology 48, 271–279.

Tierling, S., Dalbert, S., Schoppenhorst, S., Tsai, C.E., Oliger, S., Ferguson-Smith, A.C., Paulsen, M., and Walter, J. (2006). High-resolution map and imprinting analysis of the Gtl2-Dnchc1 domain on mouse chromosome 12. Genomics 87, 225–235.

Tsue, A.F., Kania, E.E., Lei, D.Q., Fields, R., McGann, C.D., Marciniak, D.M., Hershberg, E.A., Deng, X., Kihiu, M., Ong, S.E., et al. (2024). Multiomic characterization of RNA microenvironments by oligonucleotide-mediated proximity-interactome mapping. Nat Methods 21, 2058–2071.

Tucci, V. (2016). Genomic Imprinting: A New Epigenetic Perspective of Sleep Regulation. PLoS Genet 12, e1006004.

Wright, K.P., Jr., McHill, A.W., Birks, B.R., Griffin, B.R., Rusterholz, T., and Chinoy, E.D. (2013). Entrainment of the human circadian clock to the natural light-dark cycle. Curr Biol 23, 1554–1558.

Yan, J., Wang, H., Liu, Y., and Shao, C. (2008). Analysis of gene regulatory networks in the mammalian circadian rhythm. PLoS Comput Biol 4, e1000193.

Zhang, R., Lahens, N.F., Ballance, H.I., Hughes, M.E., and Hogenesch, J.B. (2014). A circadian gene expression atlas in mammals: implications for biology and medicine. Proceedings of the National Academy of Sciences of the United States of America 111, 16219–16224.

