## Supplementary material for "Light and sex modify *Snord116* genotype effects on metabolism, behavior, and imprinted gene networks following circadian entrainment": Figure S1-S6

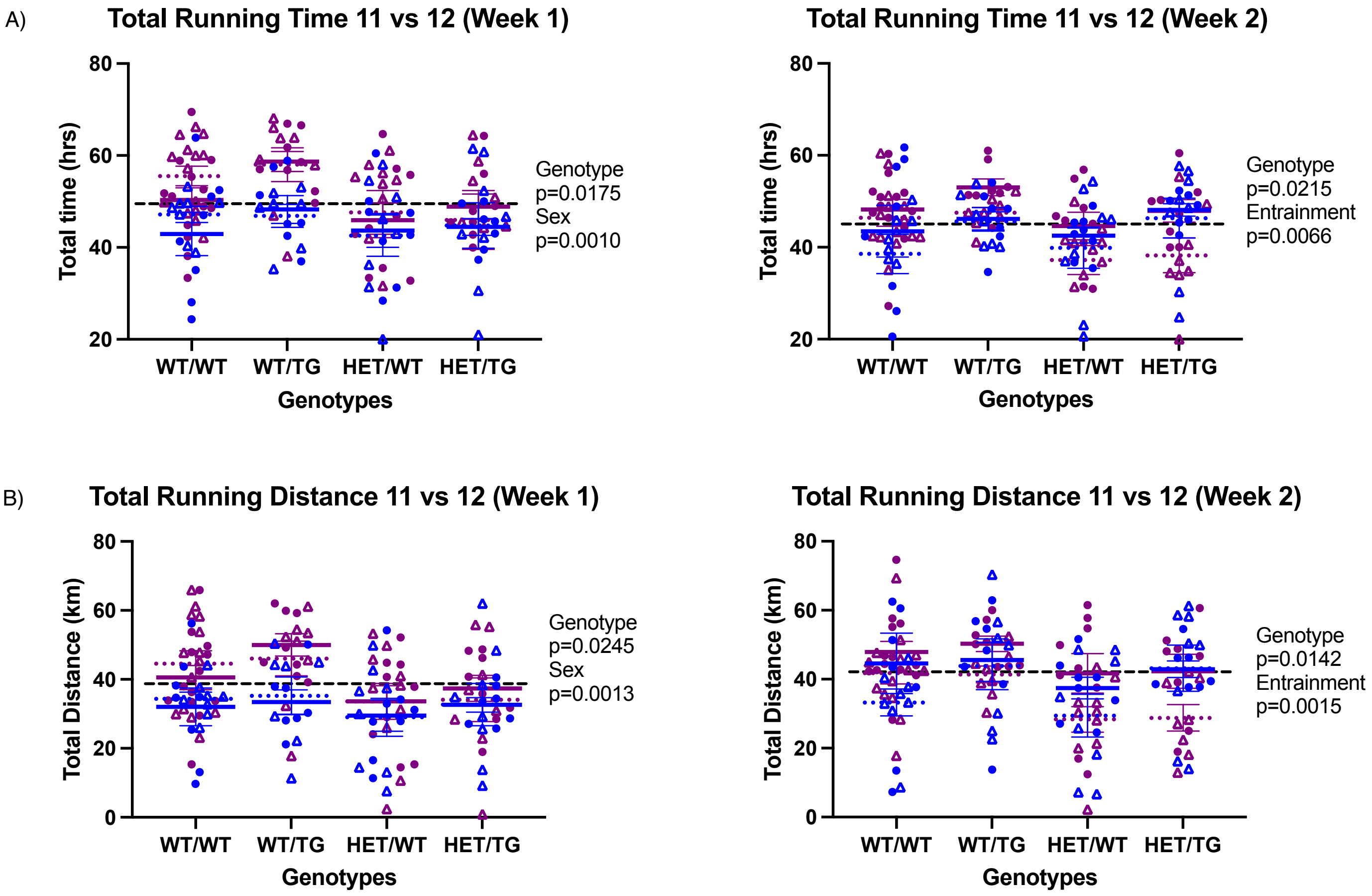


**Figure S1. Female-specific effects on total running duration and distance.** (**A**) Scatter plot of total running time measurements for each genotype and sex taken at week 1 and week 2 with 3-way ANOVA statistics of entrainment, genotype, and sex. (**B**) Scatter plot of total running distance measurements for each genotype and sex taken at week 1 and week 2 with 3-way ANOVA statistics of entrainment, genotype, and sex. Open symbols, 11:11; closed symbols, 12:12; blue symbols, male; purple symbols, female.

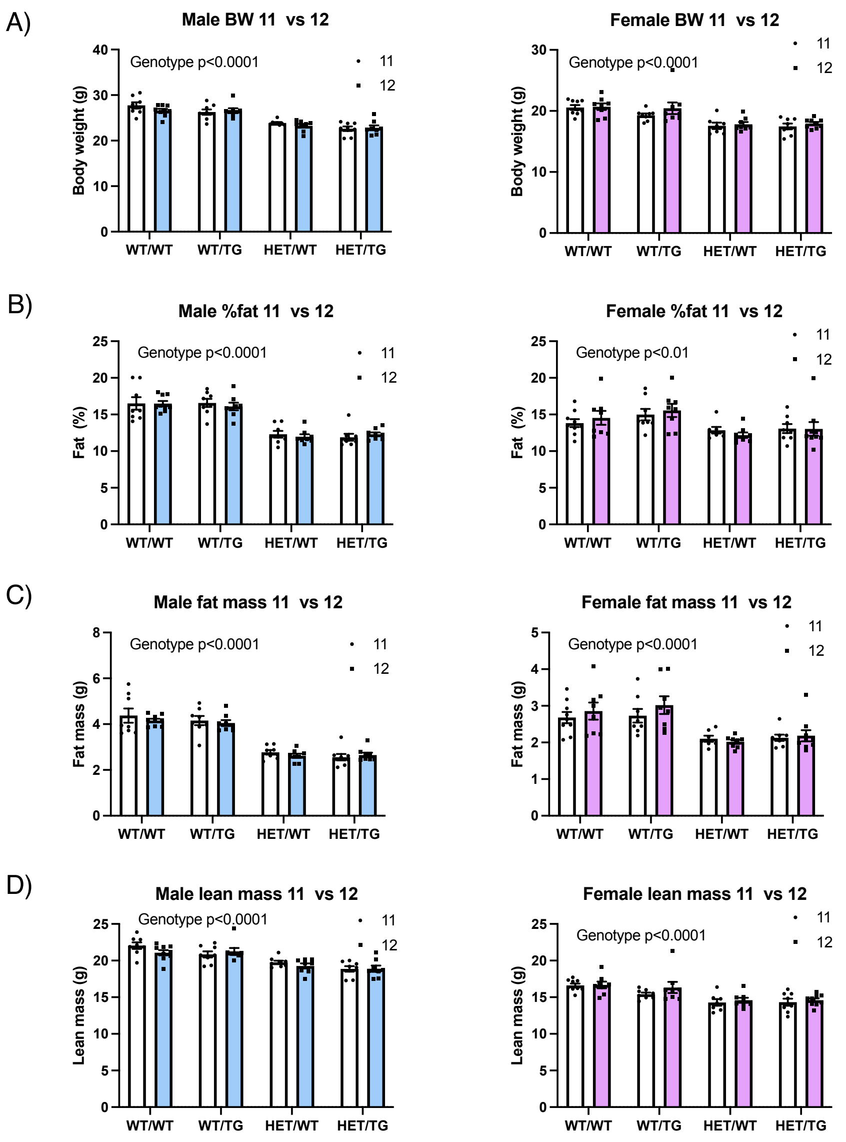


E)

| | MALES | | | FEMALES | | |
|---|---|---|---|---|---|---|
| | 2-way Anova | | | 2-way Anova | | |
| **Body Comp** | Genotype | Entrainment | Interaction | Genotype | Entrainment | Interaction |
| Body Weight | **0.0001** | 0.3699 | 0.3917 | **0.0001** | 0.1680 | 0.7555 |
| %fat | **0.0001** | 0.7523 | 0.8206 | **0.0025** | 0.7889 | 0.7826 |
| Fat mass | **0.0001** | 0.4018 | 0.7787 | **0.0001** | 0.3526 | 0.7138 |
| Lean mass | **0.0001** | 0.3576 | 0.3096 | **0.0001** | 0.1992 | 0.7709 |
| BMD | **0.0001** | 0.5610 | 0.1927 | **0.0001** | 0.6030 | 0.9174 |
| BMC | **0.0007** | 0.6440 | 0.4309 | **0.0001** | 0.7501 | 0.3975 |

**Figure S2. *Snord116* transgene does not rescue the reduced body weight phenotype in *Snord116* deletion mice.** (**A**) Body weight (BW). (**B**) % fat. (**C**) Fat mass. (**D**) Lean mass measurements taken for each genotype and sex separated by entrainment 11 (white) and 12 (skyblue – males, pink – females). (**E**) 2-way ANOVA statistics table of genotype, entrainment, and interaction effects of DEXA measurements including bone mineral density (BMD) and bone mineral composition (BMC).

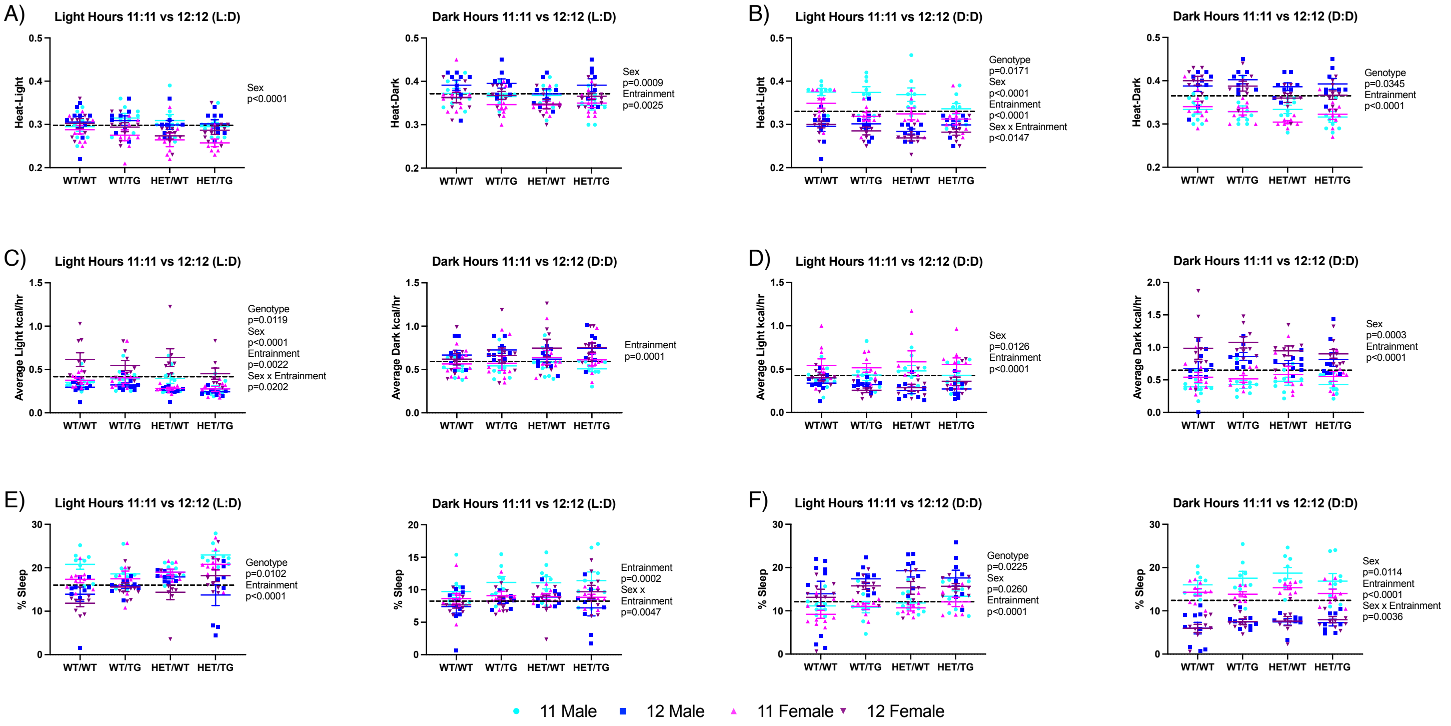


**Figure S3. Sex-specific metabolic adaptations to entrainment and *Snord116* genotype.** (**A**) L:D and (**B**) D:D scatter plots of energy expenditure (heat) measurements for each genotype and sex taken during light hours and dark hours with 3-way ANOVA statistics of entrainment, genotype, and sex. (**C**) L:D and (**D**) D:D scatter plots of food intake (kcal/hr) measurements for each genotype and sex taken during light hours and dark hours with 3-way ANOVA statistics of entrainment, genotype, and sex. (**E**) L:D and (**F**) D:D scatter plots of inactivity (% sleep) measurements for each genotype and sex taken during light hours and dark hours with 3-way ANOVA statistics of entrainment, genotype, and sex.

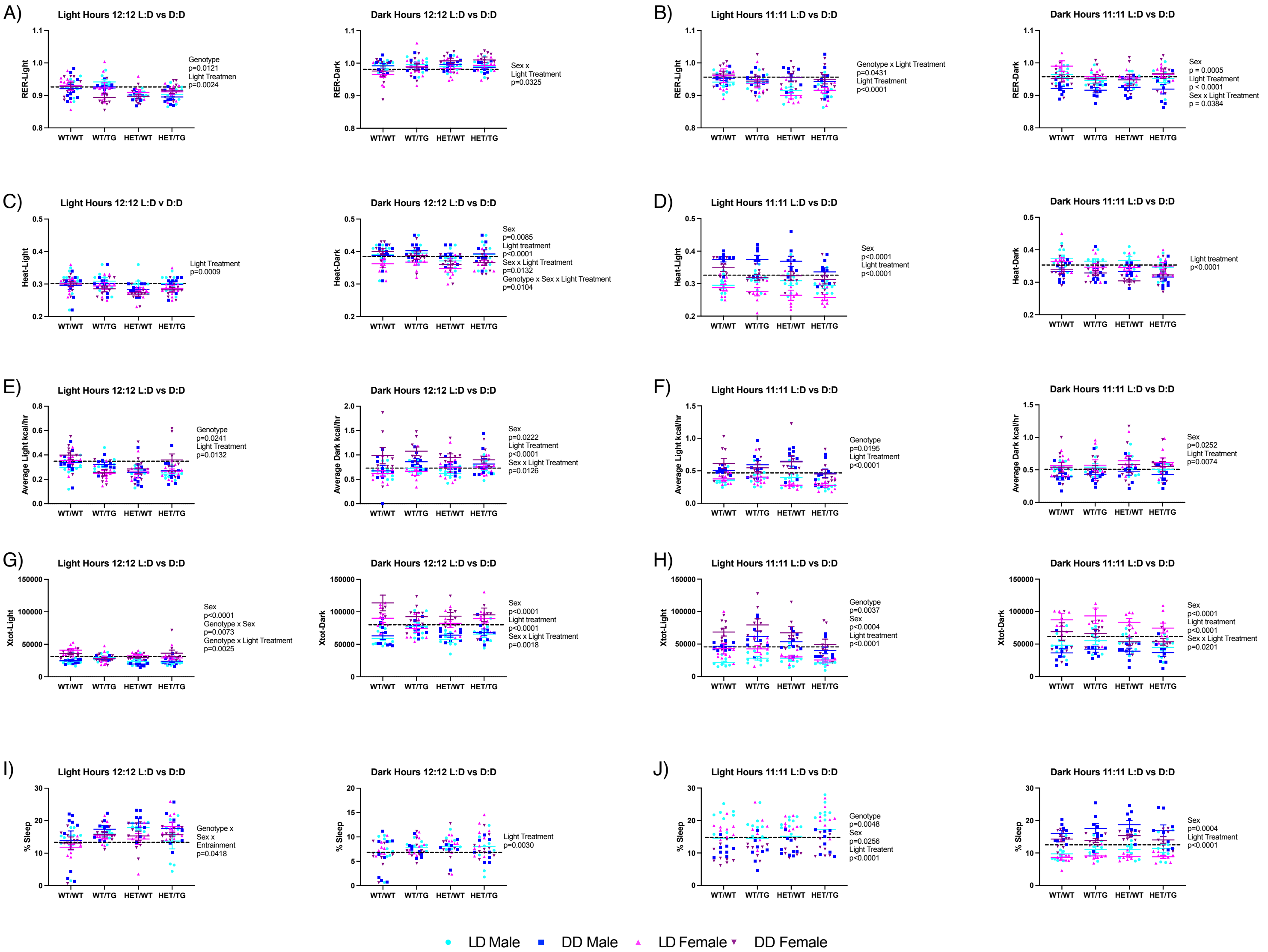


**Figure S4. Sex-specific metabolic adaptations to variable lighting conditions and *Snord116* genotype.** (**A**) 12 and (**B**) 11 scatter plots of respiratory exchange rate (RER) measurements for each genotype and sex taken during light hours and dark hours with 3-way ANOVA statistics of light treatment, genotype, and sex. (**C**) 12 and (**D**) 11 scatter plots of energy expenditure (heat) measurements for each genotype and sex taken during light hours and dark hours with 3-way ANOVA statistics of light treatment, genotype, and sex. (**E**) 12 and (**F**) 11 scatter plots of energy intake (kcal/hr) measurements for each genotype and sex taken during light hours and dark hours with 3-way ANOVA statistics of light treatment, genotype, and sex. (**G**) 12 and (**H**) 11 scatter plots of activity (X-tol) measurements for each genotype and sex taken during light hours and dark hours with 3-way ANOVA statistics of light treatment, genotype, and sex. (**I**) 12 and (**J**) 11 scatter plots of inactivity (% sleep) measurements for each genotype and sex taken during light hours and dark hours with 3-way ANOVA statistics of light treatment, genotype, and sex.

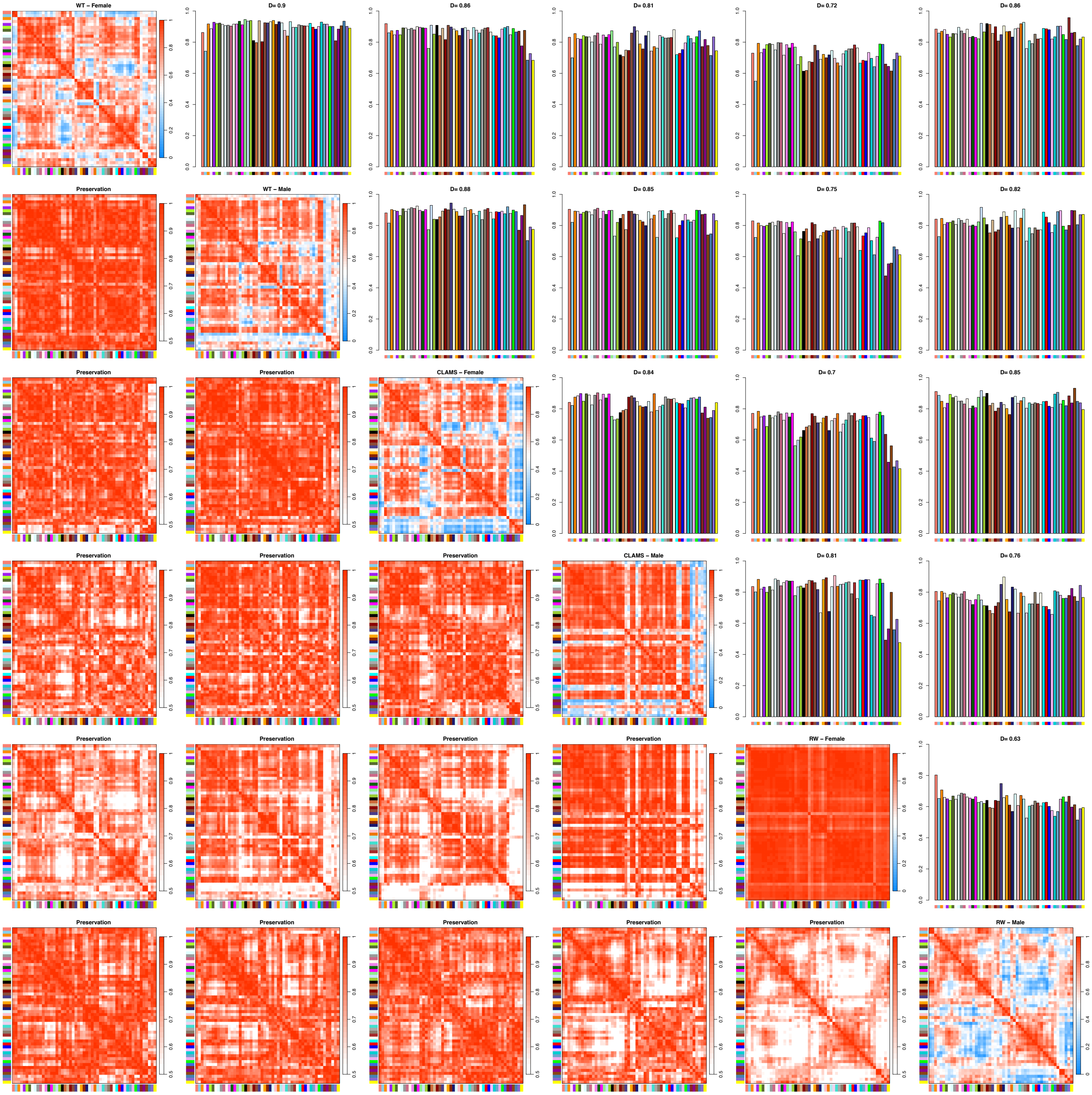


**Figure S5. WGCNA module networks are highly preserved across experiments.** Module preservation statistics of each experimental data set. Includes wild-type circadian atlas male and female, CLAMS male and female, and running wheel male and female normalized (logcpm) RNAseq counts. The density value (*D*) indicates the preservation score across all datasets. Any value *D* > 0.5 is considered highly preserved.

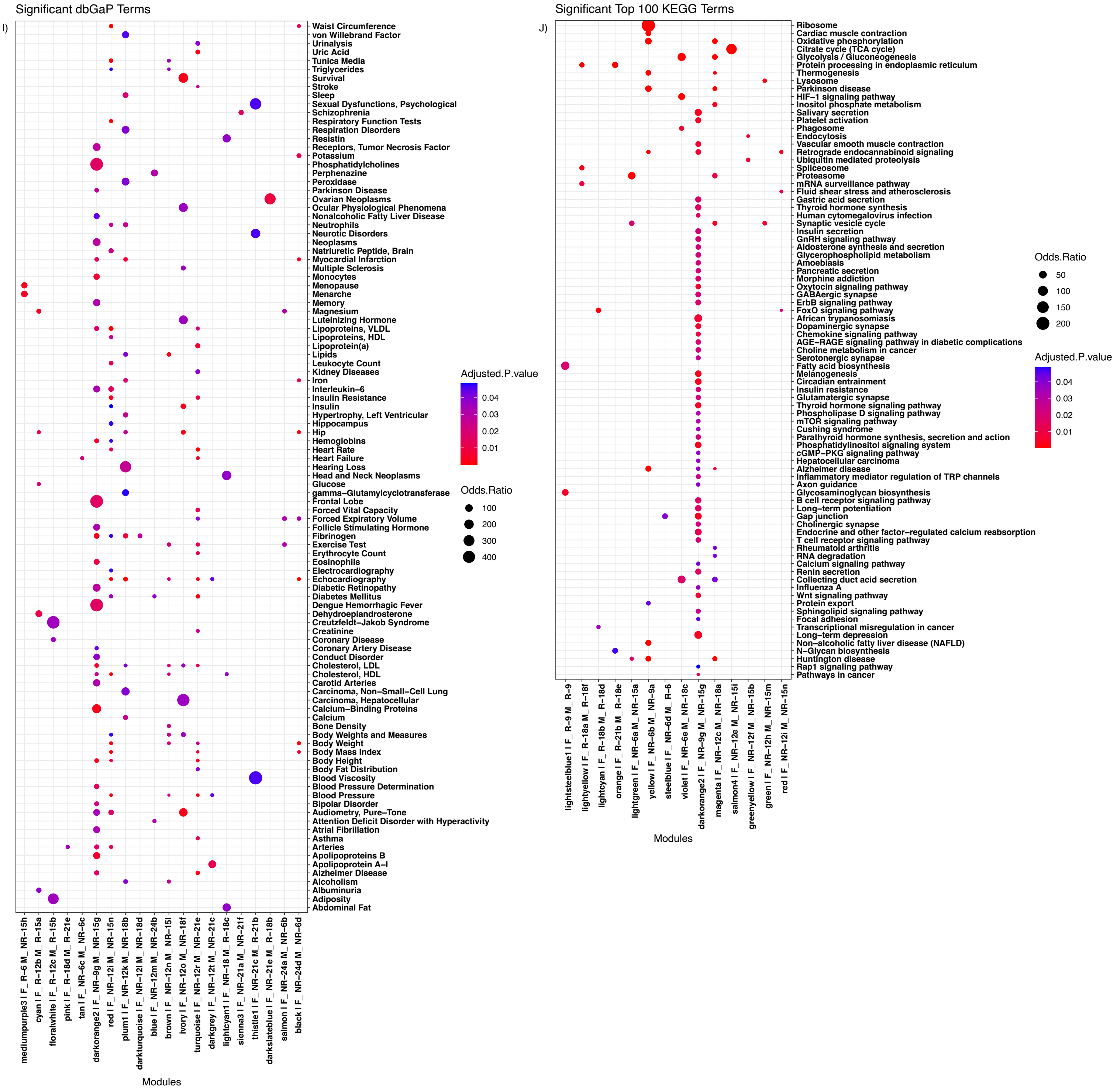


**Figure S6. Module networks highly correlated with the *Snord116* transgene are enriched for terms relevant to PWS.** (**A**) Dotplot of database of genotype and phenotypes (dbGaP) terms for modules with significant (Adjusted.pvalue < 0.05) terms. (**B**) Dotplot of top 100 KEGG for modules with significant (Adjusted.pvalue < 0.05) terms.
